# Herbaria provide a valuable resource for obtaining informative mRNA

**DOI:** 10.1101/2025.02.12.637878

**Authors:** Alexa S. Tyszka, Khong-Sam Chia, Eric C. Bretz, Linda Mansour, Drew A. Larson, Nathanael Walker-Hale, Philip Carella, Joseph F. Walker

**Affiliations:** Department of Biological Sciences, University of Illinois at Chicago, Chicago, IL 60607, USA; Cell and Developmental Biology, John Innes Centre, Norwich NR4 7UH, UK; Department of Biology, Indiana University, Bloomington, IN, 47405, USA; Department of Biosciences, Durham University, South Rd, Durham, DH1 3LE, UK

## Abstract

While DNA has built the framework for molecular insights from museum collections, the utility of archival RNA remains largely unexplored. Likely a consequence of the known instability of RNA relative to DNA, this has effectively precluded the use of herbaria for transcriptomics. Here, we challenge the assumption that herbaria cannot be used for transcriptomics by assembling transcriptomes from RNA extracted from herbarium specimens. Through systematic comparison of transcriptomes from fresh-collected, silica-dried, and archival specimens, we demonstrate the suitability of herbarium-derived RNA for transcriptomics. We show the practical applicability of archival mRNA by functionally validating a plant immune receptor synthesized from a specimen collected in 1956. These results contradict the community consensus regarding archival RNA and open the door to subsequent transcriptomic explorations of rare and extinct plant species. Our findings highlight the importance of preserving and utilizing the diversity embedded within herbarium collections.

## Introduction

The well-established field of ancient DNA (paleogenomics) provides insights into human evolution (Green *et al*., 2010), ancestral population structures of extinct organisms (Jónsson *et al*., 2014), and the evolutionary changes within crop genomes (Da Fonseca *et al*., 2015). By contrast, RNA obtained from samples stored at ambient temperatures has only recently been discussed as a source for historical insights (Friedländer and Gilbert, 2024; Tyszka *et al*., 2025).

In the context of RNA, the prefix “paleo-” and the term “ancient” have been used to refer to samples that range from hundreds to thousands of years (Smith *et al*., 2019; Mármol-Sánchez *et al.,* 2023; Mármol-Sánchez *et al.,* 2025), and the terms “historical” and “archival” have been used to discuss RNA obtained from museum specimens (Speer *et al.,* 2022; Mármol-Sánchez *et al.,* 2023; Tyszka *et al.,* 2025). To date, only a few studies have investigated historical RNA obtained from archival eukaryotic specimens. Mammalian samples have been the primary focus of these transcriptomic studies, providing important insights into the stability of mRNA (Smith *et al*., 2019; Mármol-Sánchez *et al*., 2023) and non-coding RNAs such as miRNA (Keller *et al*., 2017; Shaw *et al*., 2019; Fromm *et al*., 2021). Emerging evidence suggests that plant leaf tissue stored at room temperature may provide informative RNA, and herbaria could be a potential source of such tissue.

Viable RNA has been extracted from seeds stored for years at ambient temperatures (Rollo, 1985; Venanzi and Rollo, 1990; Fordyce *et al*., 2013; Smith *et al*., 2014; Smith *et al.,* 2017). The germination of 2000-year-old Judean date palm seeds found in desert ruins further illustrates RNA’s persistence under sub-optimal conditions (Sallon *et al*., 2008; Gros-Balthazard *et al*., 2021). Recent work has demonstrated that silica-dried leaves can retain mRNA suitable for evolutionary studies, whether the tissue is frozen after drying (He *et al*., 2022) or stored at room temperature for six months (Ruiz-Vargas *et al*., 2024). Within archival plants, viral RNA has been recovered from samples ranging from tens (Rieux *et al.,* 2021; Hartung *et al.,* 2015) to hundreds (Smith *et al.,* 2014) to even thousands of years old (Castello *et al.,* 1999). Such stability has enabled researchers to uncover the origins and timings of a plant disease emergence (Fraile *et al.,* 1997). These findings show that a variety of RNAs may be sequenced from samples stored in suboptimal conditions.

Obtaining RNA from herbaria has the potential to complement the field of plant paleogenomics by providing critical insights into gene regulation. While DNA provides a gene sequence, it cannot provide the context of when, where, and how a gene is expressed— insights that can be gained from transcriptomics. In modern genomics, sequencing a transcriptome is a central part of genome annotation and provides empirical evidence of gene expression. Historical transcriptomics, therefore, has the potential to provide an empirical basis for gene annotations for studies involving extinct or difficult-to-obtain plant samples. Improved annotations and the discovery of novel isoforms will further our understanding of the genetic variation that exists within archival samples. However, to realize the full potential of herbaria as a source of historical RNA, it is essential to assess the reliability of archival plant RNA.

Here, we demonstrate that herbaria are a promising source of informative mRNA by sequencing and assembling transcriptomes from samples dating back to 1950. Through a series of bioinformatic tests, we confirmed the reliability of the sequence data and the quality of the assembled transcriptomes. To further demonstrate the reliability of the data and provide a practical application of this approach, we identified immunity-related genes. We validated the functionality of a gene candidate from an herbarium specimen by expressing it in *Nicotiana benthamiana*. While building upon the emerging field of historical transcriptomics, this work provides a promising avenue for using plant collections to identify lost genetic variation that could be reintroduced to extant species. Furthermore, this work opens new dimensions of utility for specimens, reinforcing the importance of maintaining the world’s herbaria (Davis, 2024).

## Results

### RNA recovery, quality and sequencing results

We extracted RNA from three herbarium specimens (“-Herbarium” samples) whose vouchers are stored at MICH herbarium, with collection dates ranging from 1950 to 1995. We also investigated the quality differences between samples sequenced during the traditionally recommended timeframe (“-Fresh” samples) and their respective five-year-old silica-dried vouchers (“-Silica” samples), which were newly sequenced for this study (**Supplemental Table 1**). The RNA integrity (RIN) scores of the silica-dried and herbarium samples were consistently below the recommended level for sequencing (**Figure 1A; Supplemental Table 1**), unlike those of the fresh samples (**Supplemental Table 2**). Despite low RIN values and in two cases (*Antistrophe*-Herbarium and *Clavija*-Herbarium) RNA concentrations that were undetectable using a Qubit (see methods), RNA-seq libraries were successfully constructed, and the expected number of raw sequence reads was recovered.

**Figure 1.**
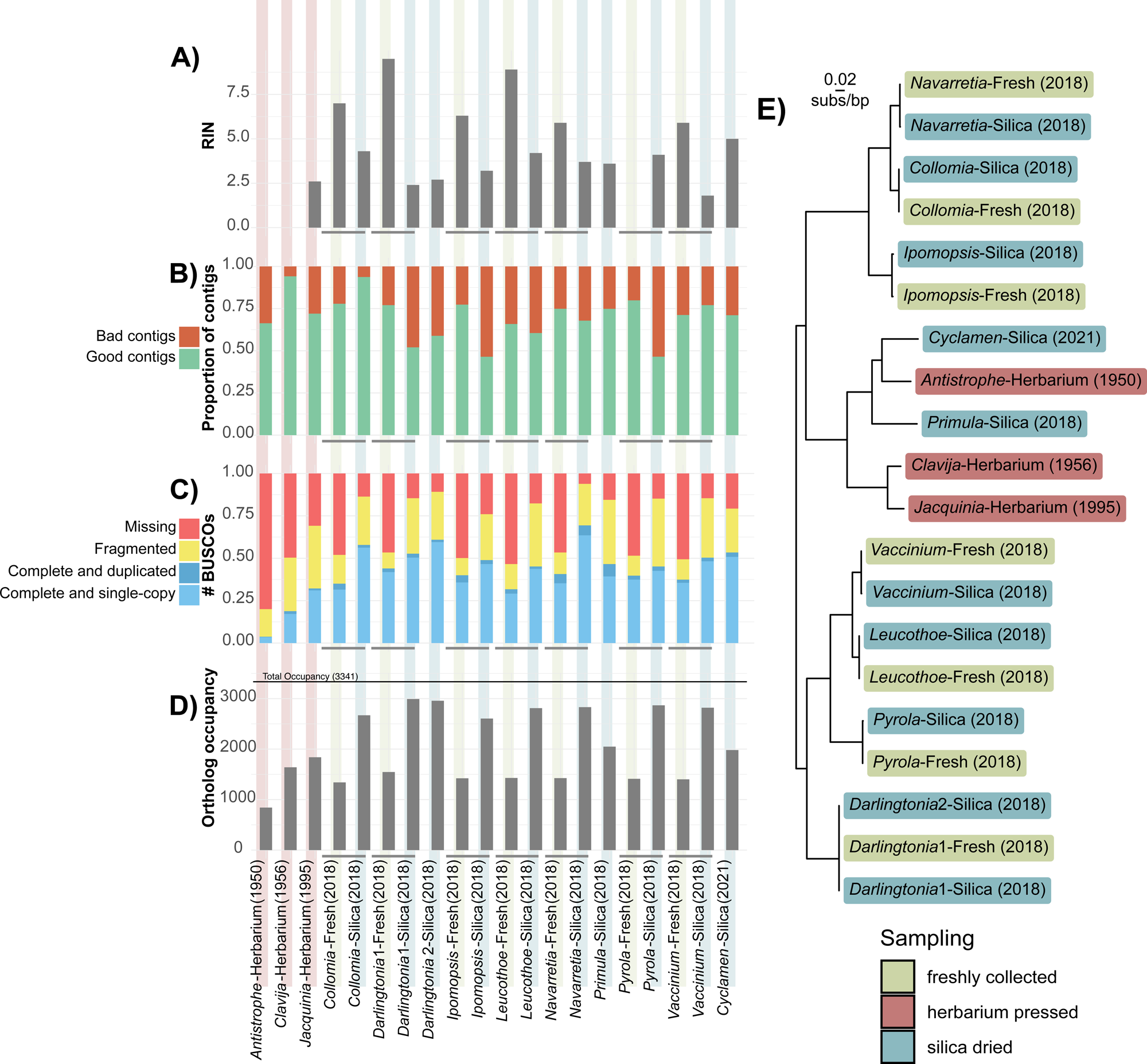
Extracted RNA from fresh, silica-dried, and herbarium specimens provide usable transcriptomes based on bioinformatic metrics. (A) The RNA integrity (RIN) scores for samples used in this study. (B) The proportion of contigs inferred to be “good contigs” compared to the proportion inferred as “bad contigs” based upon the TransRate metrics. (C) The inferred completeness of the transcriptomes based upon sequenced single copy orthologs. (D) The total number of inferred orthologs from each sample used to infer the phylogeny. (E) The coalescent-based inferred phylogenetic relationships and sample source for transcriptomes in this study. Branch lengths converted to molecular rates (subs/bp).

To ensure uniformity in the assembly procedure, the raw reads obtained from Carruthers *et al*. (2024) and the newly sequenced raw reads were all assembled using the same bioinformatics pipeline. One assessment of RNA quality we assessed was the ratio of paired to unpaired reads. Theoretically, fewer paired reads should be present as sequences degrade. When comparing results from samples collected from the same plant, the silica-dried samples had a higher proportion of unpaired sequence reads after trimming, ranging from 26.49% (*Navarretia*-Silica) to 43.69% (*Collomia*-Silica) compared to their respective fresh sample: 15.52% (*Navarretia*-Fresh) to 15.79% (*Collomia*-Fresh) (**Supplemental Table 3**). Herbarium samples had an even greater proportion of unpaired reads after trimming, with values ranging from 70.08% (*Clavija*-Herbarium) to 79.95% (*Antistrophe*-Herbarium). The loss of read pairing likely reflects a decrease in RNA quality over time, which was also captured by the shifted read-length distribution of the remaining reads that lost their pair (**Supplemental Table 4**).

To further investigate signatures of RNA degradation, we examined the length of the insert size for the fragments before sequencing. We found that the insert length for the herbarium samples was shorter than that of the fresh and silica-dried samples (**Figure 2A**; **Supplemental Fig. 1**). Finally, to test for age-related damage, we analyzed the raw sequencing data for signs of deamination. None of the samples, regardless of preservation approach, exhibited typical damage patterns (**Figure 2B; Supplemental Fig. 2**). The maximum level of adenosine to inosine (A>I) degradation was 0.12% in *Primula*-Silica, and the maximum for cytidine to uridine (C>U) was 0.13% in *Ipomopsis*-Silica.

**Figure 2.**
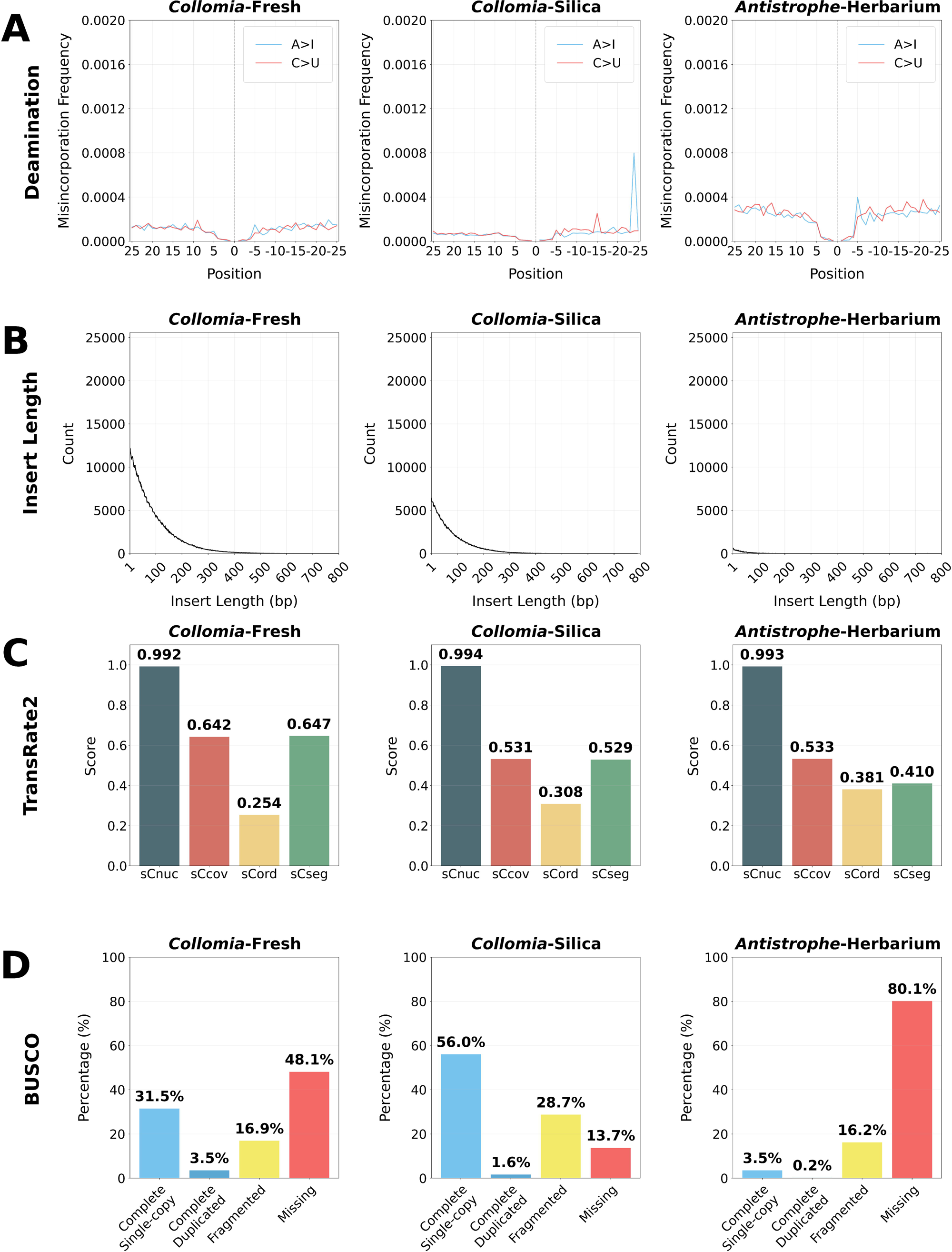
Quality assessment of sequenced and assembled RNA from representative samples for each preservation method. The quality analysis results for one representative from each preservation treatment are shown here—results for all samples are available in **Supplementary Figures 1-3**. (A) Deamination levels for adenosine to inosine (A>I) and cytidine to uridine (C>U). (B) The length of the insert for the sequenced fragments. (C) The scores for Transrate2 broken into different predicted error types: reliability of the base pair calls (*sCnuc*), the coverage level of the transcript (*sCov*), the pairing of the mapped reads (*sCord*); and evidence of chimerism within the transcripts (*sCseg*). (D) The BUSCO scores for each sample.

### Quality transcripts are obtainable from archival RNA

The quality of the assembled transcripts was assessed using the TransRate algorithm (Smith-Unna *et al*., 2016) as implemented in TransRate2 (see methods). Transrate2 is a tool designed to evaluate the quality of de-novo transcriptome assemblies, expanding on TransRate by implementing the splice-aware HISAT2 mapping approach (Kim *et al*., 2019). All transcripts that do not exhibit errors in assembly were considered “good quality”, while those with issues that exceed the allowable threshold (see methods) were considered “bad quality” (**Figure 1B)**. In brief, contig quality is inferred by analyzing common assembly errors (Smith-Unna *et al*., 2016). Between 65.8% (*Leucothoe*-Fresh) and 79.9% (*Pyrola*-Fresh) of the transcripts from the fresh tissue were regarded as good quality by this metric. For the silica-preserved samples, these numbers varied from 46.4% (*Ipomopsis*-Silica) to 93.7% (*Collomia*-Silica), and for the herbarium samples, they ranged from 66.3% (*Antistrophe*-Herbarium) to 94.1% (*Clavija*-Herbarium; **Supplemental Table 5**). Five of the seven fresh samples produced a higher proportion of good quality transcripts than their silica-dried counterpart. However, overall quality determinations and individual quality metrics were similar across preservation methods (**Supplemental Fig. 3)**. TransRate2 was also used to calculate the proportion of bases covered across each transcriptome (**Supplemental Table 5**). This value for fresh tissue ranged from 97.70-98.27%, for silica-dried tissue ranged from 93.85-96.58%, and for herbaria tissue ranged from 95.58%-97.00%. We investigated the proportion of bases covered on a contig-by-contig basis and found 100% of the bases covered ranged from 69.99%- 76.23% for fresh, 48.27%-66.92% for silica-dried and 59.05%-64.59% for herbarium samples. Contigs that did not have 100% coverage in general had over 95% coverage (**Supplemental Fig. 4**).

We examined transcriptome completeness using Benchmarking Universal Single-Copy Orthologs (BUSCOs; Simão *et al*. 2015). In all comparisons between silica-dried and fresh samples, the silica-dried samples had fewer missing BUSCOs (**Figure 1C; Supplemental Table 6**). The sequencing depth for the silica-dried samples was, on average, much greater than for fresh-collected samples (**Supplemental Table 3**). Thus, sequencing depth likely explains the greater number of complete BUSCOs recovered from silica-dried samples. These results indicate that single-copy orthologs were still identifiable in silica-dried samples, despite being stored at room temperature for five years. For transcriptomes derived from herbarium specimens, we found that the 1950s samples had fewer complete BUSCOs despite similar sequencing depth to the silica-dried samples and higher sequencing depth than the fresh samples. Missing BUSCOs ranged from 198-227 for fresh-, 26-102 for silica-, and 131-340 for herbarium-derived transcriptomes. However, two of the three counts of missing BUSCOs for herbarium samples were still within the range of those for fresh-collected samples (counts of 131 and 211 missing BUSCOs), highlighting the importance of sequencing depth when assembling transcriptomes from herbarium specimens. We fitted a multinomial logistic regression of BUSCO recovery as a function of preservation method and sequencing depth (**Supplemental Fig. 5**). As expected, proportions of complete loci increased as a function of sequencing depth while proportions of fragmented and missing loci decreased. Interestingly, the fresh samples in this analysis showed a marginally lower proportion of complete BUSCOs than silica-dried samples, and showed a lower proportion of fragmented loci.

### Archival RNA may be used for phylotranscriptomic analyses

Using all samples in the study, we inferred gene trees and a coalescent-based species tree using a phylotranscriptomic approach with the branch lengths converted to molecular rates (see methods). In total, 3,341 orthologs were inferred which contained at least 10 taxa (**Supplemental Table 7**). The lowest matrix occupancy was for *Antistrophe*-Herbarium with 842 orthologs recovered. The highest occupancy was for *Darlingtonia*1-Silica with 2,992 orthologs recovered (**Figure 1D**). The overall topology of the inferred species tree matched the current consensus relationships among species (Larson *et al*., 2020; Zuntini *et al*., 2024), and samples collected from the same plant were always sister in the tree (**Figure 1E**).

We found no evidence of longer branches subtending silica-dried or herbarium specimens, as is sometimes observed for degraded museum samples due to damage to nucleic acid damage not completely accounted for during analysis (Derkarabetian *et al*., 2019). This suggests that damaged nucleic acids did not erroneously increase branch lengths in our study.

Although assemblies from the same individual were not identical, this is not unexpected as stochastic differences introduced during library prep, sequencing, and assembly processes can result in differences in the final assembled transcript, especially for transcripts containing heterozygous sites. Importantly, this demonstrates that mRNA sequencing of herbarium samples can be used to infer evolutionary relationships among species.

### Archival RNA uncovers genes associated with plant immune systems

After we confirmed the quality of the assembled transcriptomes, we examined the NLR (Nucleotide-binding and leucine-rich repeat) immune receptor repertoires in transcriptomes from each of the three collection types (i.e., fresh, silica-dried, and herbarium samples). We selected this gene family since (i) NLRs are fast evolving and complex in architecture, (ii) their expression often correlates with function, and (iii) herbarium samples represent an untapped resource to interrogate NLR diversity. Using the NLRtracker algorithm (Kourelis *et al*., 2021), we first defined the total set of expressed NLRs in transcriptomes from fresh, silica-dried, and herbarium samples, which revealed an overall increase in detectable NLRs in silica-dried samples relative to freshly collected controls for each species (**Figure 3A, Supplemental Table 8**), potentially due to differences in sequencing depth (**Supplemental Table 3**). By comparison, fewer NLR-related genes were detected for archival RNA from the 1950s, though a full-length receptor was recovered (**Figure 3A**). Next, we generated a maximum likelihood phylogeny using the conserved NB-ARC domain to examine NLR relationships among all samples relative to a reference set of functionally validated receptors (Kourelis *et al*., 2021). Consistent with known NLR phylogenetic relationships (Chia *et al*., 2024), we observed three distinct clades corresponding to NLR subtype as defined by their N-terminal RPW8 (resistance to powdery mildew 8), TIR (toll/Interleukin-1 receptor), or CC (coiled coil) domains (**Figure 3B**). Importantly, these analyses revealed overlap between NLRs from each of the three collection types, highlighting the diversity of NLRs present within archival RNA and providing new candidates for functional investigation.

**Figure 3.**
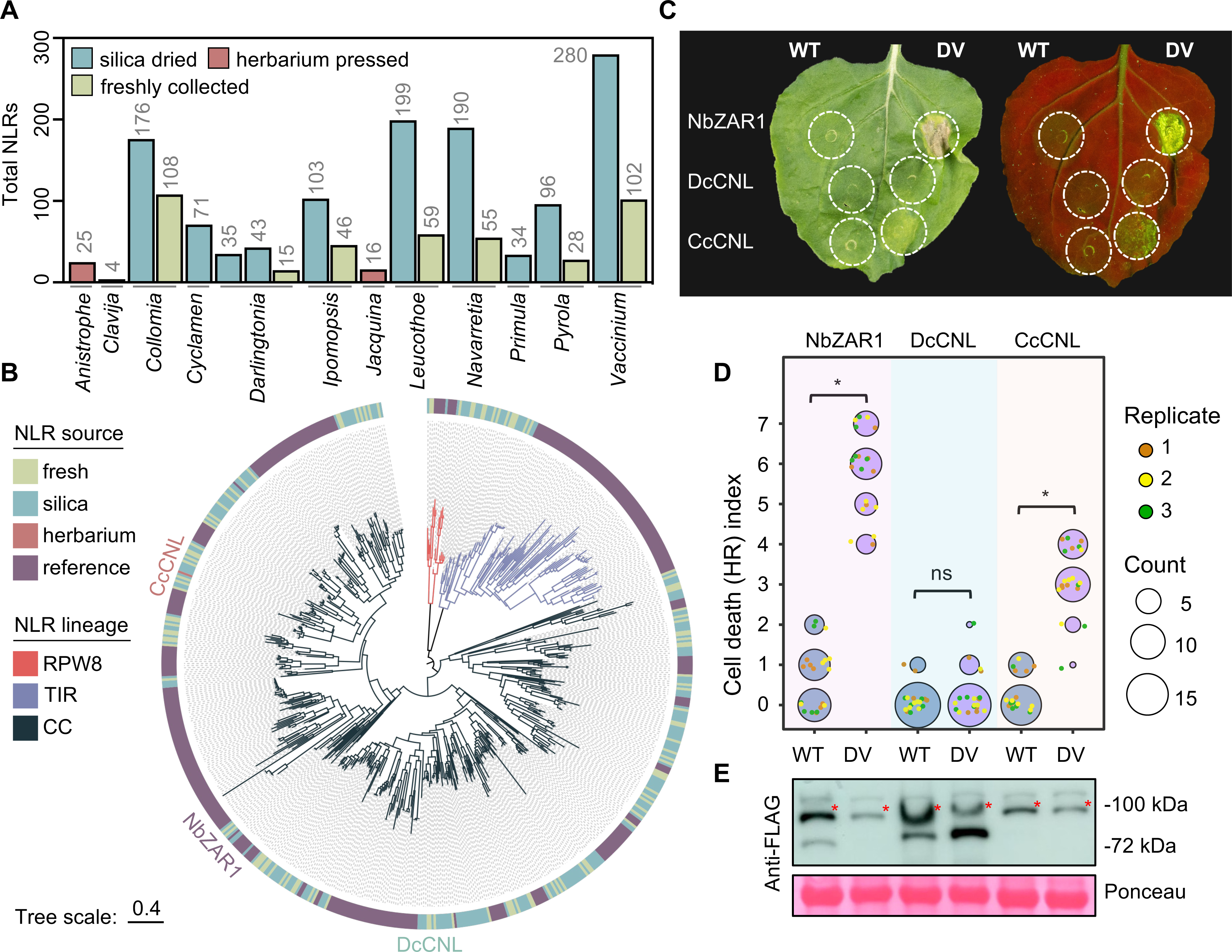
An NLR immune receptor sourced from a 68-year-old herbarium specimen of *Clavija costaricana* is functionally transferable to *Nicotiana benthamiana*. (A) Total number of NLRs identified in each transcriptome sample. (B) Maximum likelihood phylogeny of NLR immune receptors based on the central NB-ARC regulatory domain. The outer ring indicates whether each NLR was sourced from herbaria samples, silica samples, fresh samples, or belongs to a reference set of functional NLRs from flowering plants. Branch color represents NLR lineage; RPW8-NLRs (RPW8), TIR-NLRs (TIR), or CC-NLRs (CC). The three tested CC-NLRs (CNLs) are labeled on the tree. Tree scale = substitutions/site. (C) HR cell death phenotypes induced by wild-type (WT) or autoactive (DV) variants of DcCNL-3xFLAG (derived from silica-dried *Darlingtonia*), CcCNL-3xFLAG (derived from herbarium-pressed *Clavija*), or the NbZAR1-3xFLAG control in *Nicotiana benthamiana.* Leaves were imaged 7 days post agroinfiltration under bright light as well as UV light, where cell death responses appear as green fluorescence. (D) Quantification of HR cell death performed 7 d post agroinfiltration. Data from 3 independent experimental replicates are shown (n ≥ 5 infiltrations per replicate). Statistically significant differences between the means of WT and DV treatments are indicated with an asterisk * (Kruskal Wallis test, p < 0.01; ns = not significant). (E) Immunoblots of all tested NLRs 2 d post agroinfiltration. Total protein levels were visualized with Ponceau staining. Immunoblotting was repeated twice with similar results. Red asterisks (*) indicate proteins of interest.

### NLR genes recovered from archival RNA retain function

To determine whether NLRs from archival RNA retain functions in immunity, we synthesized two full-length CC-NLRs tagged with 3xFLAG epitopes for comparison against the well-established NbZAR1 CC-NLR from *Nicotiana benthamiana*. The CC-NLRs were DcCNL from a *Darlingtonia californica* sample dried with silica in 2018, and CcCNL from a *Clavija costaricana* herbarium specimen pressed in 1956. To assess their function, we synthesized and compared wild-type versions of each receptor against ‘autoactivated’ mutants carrying D (aspartic acid) to V (valine) mutations in the conserved ‘MHD’ motif known to sensitize CC-NLRs and produce immune-related cell death (Adachi *et al*., 2019). When transiently expressed in *N. benthamiana* leaves, autoactivated NbZAR1^DV^ and CcCNL^DV^ produced appreciable immune-related cell death phenotypes observable under typical light and UV illumination, whereas DcCNL^DV^ and all wild-type (WT) controls were inactive (**Figure 3CD**). Importantly, immunoblotting indicated sufficient protein accumulation for each construct in *N. benthamiana* (**Figure 3E**). Our results demonstrate that the CcCNL receptor sequence, collected in 1956 and archived at the University of Michigan Herbarium (MICH), is indeed viable and supports the idea that herbaria can be used to explore functionally relevant NLR immune receptors.

## Discussion

### Herbarium preserved tissue as a data source for coding sequence data

In this study, we present the first demonstration that plant mRNA can be obtained from herbarium specimens. Importantly, this work builds on the small but growing body of knowledge on the stability of mRNA in eukaryotic tissues (Fordyce *et al*., 2013; Smith *et al*., 2014; Smith *et al*., 2017; Smith *et al*., 2019; Mármol-Sánchez *et al*., 2023). These studies collectively overturn the preconceived notion that mRNA degrades too fast to be usable for transcriptomics.

The expanding field of historical transcriptomics can begin to provide a source of gene annotation to complement paleogenomics and supply empirical evidence about when and where particular genes were/are expressed. Unlocking the transcriptome of extinct, rare, and/or endangered plants from the convenience of an herbarium can facilitate the discovery of novel genetic variation not evident in the genome sequence alone, such as isoform production.

Although this study is limited to data from a single tissue, previous work in mammals has shown that some tissue-specific expression is maintained post-mortem (Smith *et al*., 2019; Mármol-Sánchez *et al*., 2023; Mármol-Sánchez *et al*., 2025), and future work with herbaria tissue will benefit from similar investigations.

### Sequence data shows decreased insert length but not the characteristic deamination seen in previous studies

To avoid cross-contamination, this study performed the extractions on three different days, helping ensure the RNA obtained was from the respective sample. The low RIN scores recovered for many samples (**Supplemental Table 1**, **Supplemental Document 1**) suggest that extracted RNA of low predicted quality can provide reliable transcriptomes. The lack of detectable RNA in two samples that were successfully sequenced has also been recently demonstrated for studies on the woolly mammoth (Mármol-Sánchez *et al*., 2025). Regardless, libraries were successfully prepared following the manufacturer’s protocols. In the libraries, small amounts of DNA were detected in the RNA samples despite DNase treatment. DNA is almost always present in plant samples, even when DNase treatments are performed (for example, Morales-Briones *et al*. (2021), who retained enough DNA to assemble a complete plastome despite a DNase treatment). Furthermore, the recent work on the Mammoth transcriptome showed RNA was extracted despite not using a DNase treatment (Mármol-Sánchez *et al*., 2025). To avoid cross-contamination, extractions in this study were performed at three different time points, which, in concordance with the phylogenetic structure matching expectations based on previous work in Ericales (**Figure 1E**), provides evidence for the reliability of the transcriptomic data.

We examined the sequences for the characteristic patterns of damage often seen in studies of unpreserved mRNA (Smith *et al*., 2019; Mármol-Sánchez *et al*., 2023). Read pairing loss was more prevalent in the herbarium samples than in the silica-dried or fresh ones. Loss of read pairs could occur when a shorter insert size causes the sequencer to read into adapters. Information would then be lost as adapters are trimmed during bioinformatic processing. An examination of insertion length, calculated based on the distance between the paired end reads, revealed the expected pattern: the overall insert length shortened with increasing sample age, with fresh tissue exhibiting the longest insert size and herbarium tissue the shortest. Read pairing loss and insert length provide complementary evidence that the sequences shortened, and future work that examines the rate at which this decreases and whether it influences expression patterns in a biologically meaningful way will help researchers better understand the potential insights historical transcriptomics may provide. As researchers examine older samples, it may be important to use modified protocols specifically designed for shorter reads, such as the approach taken by Smith *et al*. (2019).

To further assess the sequence data, we used a standard approach to examining deamination (Jónsson et al., 2013). The most common sources of deamination are cytidine to uridine (C>U), which is converted to C>T for sequencing, and adenosine to inosine (A>I) converted to A>G for sequencing, which have been detected in previous ancient RNA work (Smith *et al*., 2019; Mármol-Sánchez *et al*., 2023). In the present study, we did not find the same characteristic pattern of damage (**Figure 2A; Supplemental Fig. 2**). Although our current understanding of archival RNA does not allow for a detailed synthesis of what may cause this difference in deamination patterns, the sampling in this study was ∼70 years old, compared to the ∼130-year-old thylacine tissue obtained from museums. Furthermore, this tissue is derived from plant material, which may, for a yet-to-be-determined reason, help protect against deamination. Further work in the area of ancient RNA should help uncover the reasons for this pattern of deamination.

### Reliable transcriptome assemblies are obtainable from archival RNA

The individual transcriptomes used here were independently and uniformly assembled using many of the field-standard programs and approaches typically used for de novo transcriptomic assembly (Woodcock-Girard *et al*., 2025). Despite the loss of reads during trimming, each sample yielded over 10,000 predicted coding sequences, demonstrating that herbaria are a powerful source of data. The reliability of the inferred coding sequences was evaluated for accuracy using the TransRate procedure (Smith-Unna *et al*., 2016), which identifies errors common with misassembled contigs and therefore assesses the quality of assembled contigs (details in the methods section). The proportion of contigs considered “good” based on the overall quality assessment and scores for the individual quality metrics was similar across the samples regardless of preservation method. The proportion of bases that were covered by sequence reads showed a 1-2% increase for the fresh samples compared to the silica-dried and herbarium samples, indicating slightly better coverage of the assemblies from fresh tissue. On a per-contig basis, the samples showed that in both cases the majority of contigs had 100% of their bases covered, indicating that the assembled contigs were concordant with their sequencing reads. The main difference among preservation types was the number of BUSCOs recovered, with the herbarium samples from the 1950s providing the fewest complete BUSCOs. These comparisons, when taken together, demonstrate the reliability of the assembled transcripts assembled from all methods, but that herbarium samples provided fewer contigs for downstream analysis. It remains to be seen whether sequencing to greater depth or deploying programs specifically designed for archival RNA will produce assemblies similar to those of fresh tissue.

### Silica-preservation provides a functional alternative for projects seeking to obtain sequence data

Silica-preserved tissue stored at room temperature for up to six months has been shown to provide reliable coding sequence data for population genetic and phylotranscriptomic pursuits (Ruiz-Vargas *et al*., 2024). This study expanded upon this investigation and examined transcriptome quality after five years using a pairwise comparison. The seven specimen pairs consisted of one sample collected and sequenced using standard practices for field-collected RNA, as well as comparable tissue collected at the same time in silica gel and processed five years later. Phylogenetically, pairs always clustered together and showed almost no difference in subs/bp (**Figure 1E**). Based on our results, we found no advantage to storing the tissue in RNAlater for phylotranscriptomics as opposed to collecting in silica gel. Although this is a small sample size, transcriptomics from silica-collected samples deserves further investigation as it can facilitate collaborations with researchers who lack funding for specialized storage (Tyszka *et al*., 2025). Furthermore, the herbarium-derived samples also appeared in the expected phylogenetic position (Larson *et al*., 2020; Carruthers *et al*., 2024), providing further evidence that they were not the result of cross-contamination. They also did not exhibit abnormally long branches and, thus, appear to be suitable for phylotranscriptomic pursuits.

### Expressing an immune system gene from the 1950s

Although bioinformatic methods are powerful tools for data quality assessment, experimental validation remains an important standard. To provide a non-bioinformatic demonstration of the reliability of the obtained coding sequences, we validated them by synthesizing two CC-NLRs and quantifying their ability to produce cell death phenotypes when expressed in *N. benthamiana.* We demonstrate that an NLR immune receptor from a sample collected in the 1950s can be introduced and expressed in a living plant. This highlights that the predicted contigs are of reliable quality for molecular analyses, and that RNA extracted from herbaria is of sufficient quality to yield reliable transcriptomes for researchers interested in studying older specimens. The placement of historic NLRs in a phylogenetic context, combined with functional validation in living plants, serves as a confirmation that the transcriptomes obtained and assembled are accurate representations of genes that existed within the plants sampled. This finding also demonstrates the possibility of synthesizing genes from herbarium specimens, opening a new source of tissue for RNA extraction for researchers seeking coding sequences from older samples.

### Historical transcriptomics as a data source for inferring the tree of life

Here, we show that transcriptomes from herbarium specimens can be used to infer the tree of life. Phylogenetics is a field constantly adapting to incorporate novel sources of data. The One Thousand Transcriptomes project (One Thousand Plant Transcriptomes Initiative) generated a wealth of data that researchers have been able to build on and complement with their own datasets. Furthermore, transcriptomes generated for differential expression experiments are often made publicly available and can be repurposed for phylotranscriptomics (Wong and Peakall, 2022; Arreguin *et al*., 2025). When pursuing new phylotranscriptomic analyses, these existing data sources may be complemented with data from herbaria, so contentious and difficult-to-acquire lineages can be included in the analysis.

The predominant data source for phylogenomic analyses using herbaria has been probe datasets, which rely on DNA extraction (Johnson *et al*., 2019; Zuntini *et al*., 2024). Although probe-based DNA sequencing provides a powerful method of targeting specific genes, these methods are largely limited to the set of genes targeted *a priori,* and unless the same probe set is used, the datasets are not compatible. As probe datasets are often built using transcriptomes as a guide, the coding regions sequenced from transcriptomes can be used to complement existing probe datasets and transcriptomic data. The inclusion of transcriptomic data with probe datasets has allowed researchers to refine not only phylogenetic hypotheses, but patterns of gene and genome duplication (Lagou *et al*., 2025). The ability to obtain transcriptomes from herbaria not only has the potential to expand our knowledge of the tree of life but also to provide insights into the processes that have shaped and continue to shape it.

### Herbaria as a novel source of historical transcriptomic insights

A body of literature has slowly developed, highlighting that RNA in seeds maintains its integrity for germination under extreme conditions (Sallon *et al*., 2008; Gros-Balthazard *et al*., 2021), and this can even be leveraged for RNA extraction (Rollo, 1985; Fordyce *et al*., 2013). Ancient mummies have had their mRNA extracted (Venanzi and Rollo 1990), and canids found in permafrost have had their transcriptomes profiled (Smith *et al*., 2019). The expression profiles of museum tissue show that some signals of tissue-specific expression are maintained (Mármol-Sánchez *et al*. 2023). Furthermore, microRNAs have been extracted from a variety of eukaryotic species (Keller *et al*., 2017; Shaw *et al*., 2019; Fromm *et al*., 2021), and with these data, scientists are gaining new insights into the gene regulation of extinct organisms (Mármol-Sánchez *et al.,* 2023; Mármol-Sánchez *et al.,* 2025).

### Limitations of this study and future directions for understanding herbaria as a source of transcriptomic data

The sampling scheme and structure of the study leave several questions to be answered. Our focus on mRNA raises the question of how reliably microRNAs may also be obtained from these samples. Furthermore, in this study, we lacked replication of our samples; therefore, we cannot reliably analyze biologically meaningful differences in expression patterns. Expression patterns can help answer questions such as whether the immune response to past exposure to pathogens is still identifiable in herbarium specimens. Further, this can address whether the stress of desiccation is reflected in expression patterns. Another limitation we have is the lack of reference genomes for the samples in question. We are unable to analyze exon-exon junctions and exonic vs. intronic reads. Without replicates and complete genomes, we were unable to determine whether there is biased retention of genes and exonic regions. Further research may help answer this and other questions, such as whether different plant tissue types, such as succulent versus papery tissues, vary in how well they preserve mRNA. How does the length and type of preservation influence degradation, and at what age (and to what extent) will deamination play a role in herbarium-based transcriptomic studies? The ability to access mRNA from herbaria opens the door for these and many other questions. Here, we provide the first step by establishing that usable plant mRNA may be obtained from herbarium-sourced tissue.

In summary, our work (i) demonstrates that mRNA may be obtained from pressed herbarium specimens and years-old silica-dried samples; (ii) this mRNA may be sequenced and assembled into reliable transcriptomes that can be used for a variety of pursuits including phylotranscriptomics; and (iii) using transcriptomes from herbarium specimens, we can predict NLR genes, indicating the continued presence of genetic information including essential variation underlying disease resistance. To date, work on historical RNA remains almost nonexistent across the tree of eukaryotes, with great potential for insights across broad disciplines in biology (Friedländer and Gilbert, 2024; Tyszka *et al*., 2025). This work further highlights the value of the world’s herbaria by explicitly combining the historical collections with insights from high-throughput sequencing. Our results emphasize the importance of protecting these invaluable resources as novel uses for them continue to arise.

## Methods

### Sampling Scheme

For this study, we chose to sample members of the flowering plant order Ericales, a clade of over 13,000 species that contains important crops such as tea, kiwi, and blueberries, ornamentals, parasitic lineages, and carnivorous plants (Rose *et al*., 2018; Larson *et al*., 2020; Carruthers *et al*., 2024). We used samples stored under three preservation approaches: RNAlater followed by freezing (i.e., “-Fresh” samples), silica-dried tissue collected in abundant silica gel in the field (i.e., “-Silica” samples), and herbarium tissue (i.e., “-Herbarium” samples). Our sampling included seven paired silica-dried leaves and corresponding fresh leaves, in six cases collected from the same individual plant, and in all cases at the same time and from the same tissue type. Our sampling also included three herbarium specimens collected between 1950 and 1995 and three additional silica-dried samples (**Supplemental Table 9**).

### RNA extraction

We obtained mRNA from three herbarium samples and 10 silica-dried samples, which were stored at room temperature for ca. 5 years (Supplemental Table 1). Tissue grinding, RNA extractions with the DNAse treatment, and quality assessment were performed using the same procedure as Ruiz-Vargas *et al*. (2024). To ensure the extraction procedure did not introduce cross-contamination, the extractions were performed at three different times. The silica dried samples were extracted on June 4^th,^ 2023, and June 11^th,^ 2023, and the herbarium samples were extracted on June 12^th^, 2023. For two of the samples, the RNA concentrations measured with a Qubit fluorometer (Thermo Fisher) were below the minimum concentration required for analysis on the TapeStation (**Supplemental Table 1; Supplemental Document 1**). Ribosomal RNAs were removed using the QIAseq FastSelect kit (Qiagen), and the RNA-seq libraries were prepared with the Kapa Hyper Stranded mRNA library kit (Roche). Paired-end 150 bp was carried out on a single lane of an Illumina NovaSeq X.

### Transcriptome Assembly and Quality Assessment

Transcriptomes for all samples, including the fresh samples from Carruthers *et al*. (2024), were assembled using the same procedure described below. The quality of the raw sequence reads was initially assessed using the program FastQC (https://www.bioinformatics.babraham.ac.uk/projects/fastqc/). Single base pair errors were corrected using the program Rcorrector v1.0.4 (Song and Florea, 2015), with reads deemed unfixable removed using the unfixable_filter.py script from Morales-Briones *et al*. (2021). Trimmomatic v0.39 (Bolger et al., 2014) was used to remove adapter sequences and trim low-quality reads.

Kraken2 v2.1.2 (Wood *et al*., 2019) was used to classify any material matching the standard protist and fungal database, and any classified material was removed as a contaminant. The same procedure was then used to classify reads that matched the chloroplast and then the mitochondrion database from the program Kakapo (Ramanauskas and Igić, 2023). The remaining reads were classified as nuclear reads.

Trinity v2.1.1(Haas *et al*., 2013) was used to assemble the nuclear sequence reads into transcripts. Chimeric transcripts were detected using the procedure of Yang and Smith (2013) with a database composed of the peptide sequences for *Arabidopsis thaliana* (TAIR10) and *Actinidia chinensis* (Red5_PS1_1.69.0). The abundances of the nuclear reads were then quantified using Salmon v1.3.0 (Patro *et al*., 2017) and clustered into the highest supported transcript using Corset v1.09 (Davidson and Oshlack, 2014). The open reading frames were predicted using TransDecoder v5.5.0 (https://github.com/TransDecoder/TransDecoder), and the best supported ORF was retained. In total, we generated 13 new transcriptomes, 3 of which originated from herbarium material, and reassembled 7 previously sequenced transcriptomes.

The completeness of the assemblies was assessed using BUSCO v.5.8.2 (Manni *et al*., 2021) with the “--mode transcriptomes” option and the viridiplantae_odb10 database. The quality of the transcripts was assessed using TransRate2 (https://github.com/ericbretz/transrate2), an implementation of the TransRate procedure (Smith-Unna *et al*., 2016). TransRate2 is a tool designed to evaluate the quality of de-novo transcriptome assemblies, expanding on TransRate by implementing the splice-aware HISAT2 mapping approach (Kim *et al*., 2019). Specifically, the score a transcript is penalized based on the following criteria: (i) The transcriptome shows evidence of chimeric sequences, (ii) The transcript has low coverage from the raw reads, (iii) The reads mapping to the transcript are not properly paired and (iv) A base in the transcript does not correspond to the base of the sequences that are mapped to it. Given sufficient penalties, the transcript will be considered “bad quality”; otherwise, transcripts will be considered “good quality”. The mapping rates obtained from TransRate2 were used to calculate the total coverage across all bases and the per-contig base coverage.

### RNA Damage Analysis

To determine the length of the insert size associated with each read pair, we used HISAT2 to map the cleaned raw reads back to the assembled transcriptome. The output BAM file was then used to calculate the distance between the reads using a novel Python script (https://github.com/ericbretz/Herbaria/). The program mapDamage2 v.2.3.0a0-10bc442 (Jónsson *et al*., 2013) with “--merge-libraries” and all other settings set to default was used to test for deamination, with the same BAM file generated for the insert size analysis used as input.

### Phylogenetic Analysis

We conducted a phylogenomic analysis by identifying homologous gene clusters from the assembled transcriptomes of fresh, silica-dried, and herbarium samples. The phylogenomic analysis also included a previously assembled transcriptome of *Cornus florida* (NCBI RefSeq assembly: GCF_030987335.1). CD-Hit v4.8.1(Fu *et al*., 2012) with a sequence identity threshold of 0.99 was used to reduce redundancy in the transcriptomes. An all-by-all BLASTN (Altschul *et al*., 1990) search on the transcriptomes was conducted for initial homology detection, with an e-value of 10 and the maximum target sequences set to 1000. The homolog clusters, as detected by BLASTN, were then further refined using mcl v14-137 (Van Dongen, 2008), with an inflation value of 1.4. All remaining homolog clusters with at least 10 taxa were retained for phylogenomic analysis.

The putative homologs were aligned using MAFFT v7.490 (Katoh and Standley, 2013) with the “--auto” and “--maxiterate 1000” settings. The phyx program pxclsq v1.2 (Brown *et al*., 2017) was used to remove all alignment columns with occupancy less than 10%. Maximum likelihood, as implemented in IQ-TREE v1.6.12 (Nguyen *et al*., 2015), with the GTR model of evolution and gamma rate variation, was used to infer a phylogenetic tree for each homolog. We chose the GTR+G model because it is the most parameter-rich within the GTR family, and other commonly used time-reversible models are nested within it. From the inferred trees, all tips that had branch lengths of 0.2 subs/bp and/or were 10x longer than their sister or 0.3 subs/bp absolute value were removed, and all clades in trees that consisted of gene sequences from a single sample were collapsed to retain only the one with the most information in the alignment. The refined homologs were then run through the same procedure three more times to refine the phylogenetic dataset further.

Orthologous groups were extracted from the homolog trees using the maximum inclusion method (Yang and Smith, 2014), requiring a minimum of 10 taxa and removing any branch lengths with a relative value of 0.2 subs/bp or an absolute value of 0.3 subs/bp. The sequences corresponding to the extracted ortholog trees were then aligned using MAFFT, cleaned using pxclsq as before, and used to estimate gene trees with IQ-TREE using the same settings as the homology procedure. A species tree was then inferred using the maximum quartet support species tree method implemented in ASTRAL v5.7.8 (Zhang *et al*., 2018). The coalescent units were converted to subs/bp using the median branch length value of all concordant branches in the gene trees as implemented in the program BES (Walker *et al*., 2022).

### NLR identification and phylogenetic analysis

NLR identification was performed using the default settings of the NLRtracker pipeline, which is available online (https://github.com/slt666666/NLRtracker). Phylogenetic analysis was performed by comparing the central NB-ARC domains of fresh, silica-dried, and herbarium samples against a reference set of flowering plant NLRs (Kourelis *et al*., 2021). Isolated domains were aligned using MAFFT and trimmed using trimAl (Capella-Gutiérrez et al., 2009). The resulting alignments were used for maximum likelihood phylogenetic analysis with IQ-TREE v2.0.3 using the “JTT+F+G4” model selected by ModelFinder (Kalyaanamoorthy *et al*., 2017) and 1000 ultrafast bootstrap replicates to assess support (Hoang *et al*., 2018). The phylogenetic tree was subsequently rendered using TVbot (Xie *et al*., 2023).

### Plant growth conditions

*N. benthamiana* plants were grown in soil under controlled conditions, with a long-day photoperiod (16 h light; 160–200 μE, using Sylvania F58W/GRO fluorescent lighting) at 22°C.

### Cloning, immunoblotting, and cell death assays

Constructs for cell death assays were assembled using the Modular Cloning (MoClo) system with the following components: pICH85281 (mannopine synthase promoter + W [MasWpro], Addgene #50272), pICSL50007 (AddGene #50308), pICSL60008 (Arabidopsis heat shock protein terminator [HSPter], TSL SynBio), binary vector pICH47742 (Addgene #48001), and the appropriate synthesized NLR gene or variant. The CC-NLRs synthesized were 3xFLAG epitope tagged wild-type (WT) and autoactive (DV) variants of DcCNL (identified from the *Darlingtonia*-Silica transcriptome) and CcCNL (identified from the *Clavija*-Herbarium transcriptome). NbZAR1 constructs were generated using parts from an established goldengate toolkit (Harant *et al*., 2022). All resulting constructs were transformed into *Agrobacterium tumefaciens* GV3101 (pMP90) by electroporation. Cell death assays were performed by Agrobacterium-mediated transient expression, where fully expanded leaves of 4-week-old *N. benthamiana* plants were infiltrated with *A. tumefaciens* strains containing a binary expression plasmid mixed at a 1:1 ratio with a suspension of *A. tumefaciens* containing the p19 silencing suppressor. Bacterial cells were resuspended in a fresh infiltration buffer (10 mM MES-KOH, 10 mM MgCl₂, and 150 mM acetosyringone, pH 5.6) and adjusted to an OD600 of 0.3. Immune-related cell death phenotypes were assessed using an HR index ranging from 0 (no visible symptoms) to 7 (fully confluent cell death). For immunoblotting, protein samples were prepared from six *N. benthamiana* leaf discs (8mm diameters) at 2 d after agroinfiltration. Leaf samples were then ground to a fine powder and homogenized in 150 μL of 2x LDS sample buffer (NuPAGE LDS Sample Buffer). Immunoblotting was performed with anti-flag antibody coupled with HRP (Sigma-Aldrich A8592, 1:5,000 dilution), and total protein loading was visualized with Ponceau S solution (Sigma-Aldrich, P7170).

### Statistical analysis

We estimated the differences in recovery rate of BUSCO loci from transcriptomes derived from fresh, silica dried, and herbarium tissue, accounting for differences in sequencing depth. We fitted a multinomial logistic regression using nnet v7.3-20 (Venables and Ripley, 2002), with counts of complete (summing single copy and duplicated loci), fragmented and missing BUSCOs as a response, and storage condition and read depth as predictors. To assess the inferences of this model, we visualized the predicted proportions of recovery from each category per sequencing depth. All analyses were conducted in R v4.5.1 (R Core team, 2025).

## Data Access

All data supporting the conclusions of this study are present in the paper, supplementary material, or available publicly. The raw sequence reads generated for this study have been submitted to the NCBI BioProject database (https://www.ncbi.nlm.nih.gov/bioproject/) under accession number PRJNA1200749. Voucher information is available in **Supplemental Table 9**. Analysis outputs for TransRate2 and BUSCO are available as **Supplemental Tables 5 & 6**. Assembled sequences, trees, and NLRtracker outputs generated in this study have been submitted to Zenodo (https://zenodo.org/records/14720388).

## Acknowledgements

The authors thank the UIC Sequencing Core, Zarema Arbieva, and Nina S. Los for library prep and quality assessment, Tom Carruthers for assistance in organizing the sequence data for the fresh tissue samples, Jim Belsher-Howe and Plumas National Forest staff for logistical support and permission to collect specimens, and Brad Ruhfel for advice on the herbarium collections. Steve Kelly provided valuable insights on the function of TransRate. A.S.T. was supported by the National Science Foundation (NSF) grant GRFP-2236870. D.A.L. was supported by NSF IOS-2109716. J.F.W. acknowledges start-up funding support from the University of Illinois Chicago. P.C. was supported by UKRI (UK Research and Innovation) Biotechnology and Biological Sciences Research Council (BBSRC) core funding [BB/X010996/1]. The authors thank the University of Michigan Herbarium (MICH) for providing access to archival specimens. The authors also thank three anonymous reviewers for their helpful comments which have improved this manuscript.

## Author Contributions

A.S.T, J.F.W, D.A.L and P.C led the research design. A.S.T performed the RNA extractions. All authors contributed to conducting the research. A.S.T, E.C.B, N.W-H and P.C. prepared the figures. A.S.T, J.F.W and P.C led the writing of the paper with contributions from all authors.

## References

Adachi, H. et al. (2019) ‘An N-terminal motif in NLR immune receptors is functionally conserved across distantly related plant species’, eLife, 8, p. e49956. Available at: 10.7554/eLife.49956.

Altschul, S. et al. (1990) ‘Basic Local Alignment Search Tool’, Journal of Molecular Biology, (3), pp. 403–410. Available at: 10.1016/S0022-2836(05)80360-2.

Arreguin, Shawn, Nathanael Walker-Hale, Madeline R. Casagrande, Savannah Bishop, Eliza Pugacewicz, Mohammed Ramizuddin, Caili Savitzky et al. “Going green: Recycling transcriptomes to infer evolutionary relationships, gene duplication, gene tree conflict, and patterns of molecular evolution in the Apocynaceae.” bioRxiv (2025): 2025–06.

Bolger, A.M., Lohse, M. and Usadel, B. (2014) ‘Trimmomatic: a flexible trimmer for Illumina sequence data’, Bioinformatics, 30(15), pp. 2114–2120. Available at: 10.1093/bioinformatics/btu170.

Brown, J.W., Walker, J.F. and Smith, S.A. (2017) ‘Phyx: phylogenetic tools for unix’, Bioinformatics. Edited by J. Kelso, 33(12), pp. 1886–1888. Available at: 10.1093/bioinformatics/btx063.

Capella-Gutiérrez, S., Silla-Martínez, J.M. and Gabaldón, T. (2009) ‘trimAl: a tool for automated alignment trimming in large-scale phylogenetic analyses’, Bioinformatics, 25(15), pp. 1972–1973. Available at: 10.1093/bioinformatics/btp348.

Carruthers, T. et al. (2024) ‘Repeated shifts out of tropical climates preceded by whole genome duplication’, New Phytologist, 244(6), pp. 2561–2575. Available at: 10.1111/nph.20200.

Castello, J. D. et al. Detection of tomato mosaic tobamovirus RNA in ancient glacial ice. Polar Biol. 22, 207–212 (1999). 10.1007/s003000050411

Chia, K.-S. et al. (2024) ‘The N-terminal domains of NLR immune receptors exhibit structural and functional similarities across divergent plant lineages’, The Plant Cell, 36(7), pp. 2491–2511. Available at: 10.1093/plcell/koae113.

Da Fonseca, R.R. et al. (2015) ‘The origin and evolution of maize in the Southwestern United States’, Nature Plants, 1(1), p. 14003. Available at: 10.1038/nplants.2014.3.

Davidson, N.M. and Oshlack, A. (2014) ‘Corset: enabling differential gene expression analysis for de novoassembled transcriptomes’, Genome Biology, 15(7), p. 410. Available at: 10.1186/s13059-014-0410-6.

Davis, C.C. (2024) ‘Collections are truly priceless’, Science, 383(6687), pp. 1035–1035. Available at: 10.1126/science.ado9732.

Derkarabetian, S., Benavides, L.R. and Giribet, G. (2019) ‘Sequence capture phylogenomics of historical ethanol-preserved museum specimens: Unlocking the rest of the vault’, Molecular Ecology Resources, 19(6), pp. 1531–1544. Available at: 10.1111/1755-0998.13072.

Fordyce, S.L., et al. (2013) ‘Deep Sequencing of RNA from Ancient Maize Kernels’, PLoS ONE. Edited by D.Q. Fuller, 8(1), p. e50961. Available at: 10.1371/journal.pone.0050961.

Fraile, A. et al. A century of tobamovirus evolution in an Australian population of Nicotiana glauca. J. Virol. 71, 8316–8320 (1997).

Friedländer, M.R. and Gilbert, M.T.P. (2024) ‘How ancient RNA survives and what we can learn from it’, Nature Reviews Molecular Cell Biology, 25(6), pp. 417–418. Available at: 10.1038/s41580-024-00726-y.

Fromm, Bastian, Marcel Tarbier, Oliver Smith, Emilio Marmol-Sanchez, Love Dalen, M. Tom P. Gilbert, and Marc R. Friedländer. “Ancient microRNA profiles of 14,300-yr-old canid samples confirm taxonomic origin and provide glimpses into tissue-specific gene regulation from the Pleistocene.” RNA 27, no. 3 (2021): 324–334.

Fu, L. et al. (2012) ‘CD-HIT: accelerated for clustering the next-generation sequencing data’, Bioinformatics, 28(23), pp. 3150–3152. Available at: 10.1093/bioinformatics/bts565.

Green, R.E. et al. (2010) ‘A Draft Sequence of the Neandertal Genome’, Science, 328(5979), pp. 710–722. Available at: 10.1126/science.1188021.

Gros-Balthazard, M. et al. (2021) ‘The genomes of ancient date palms germinated from 2,000 y old seeds’, Proceedings of the National Academy of Sciences, 118(19), p. e2025337118. Available at: 10.1073/pnas.2025337118.

Haas, B.J. et al. (2013) ‘De novo transcript sequence reconstruction from RNA-seq using the Trinity platform for reference generation and analysis’, Nature Protocols, 8(8), pp. 1494–1512. Available at: 10.1038/nprot.2013.084.

Harant, Adeline, Hsuan Pai, Toshiyuki Sakai, Sophien Kamoun, and Hiroaki Adachi. “A vector system for fast-forward studies of the HOPZ-ACTIVATED RESISTANCE1 (ZAR1) resistosome in the model plant Nicotiana benthamiana.” Plant Physiology 188, no. 1 (2022): 70–80.

Hartung, J.S. et al. (2015) History and diversity of Citrus leprosis virus recorded in herbarium specimens. Phytopathology 105, 1277–1284

He, J. et al. (2022) ‘A phylotranscriptome study using silica gel-dried leaf tissues produces an updated robust phylogeny of Ranunculaceae’, Molecular Phylogenetics and Evolution, 174, p. 107545. Available at: 10.1016/j.ympev.2022.107545.

Hoang, D.T. et al. (2018) ‘UFBoot2: Improving the Ultrafast Bootstrap Approximation’, Molecular Biology and Evolution, 35(2), pp. 518–522. Available at: 10.1093/molbev/msx281.

Johnson, Matthew G., Lisa Pokorny, Steven Dodsworth, Laura R. Botigué, Robyn S. Cowan, Alison Devault, Wolf L. Eiserhardt et al. “A universal probe set for targeted sequencing of 353 nuclear genes from any flowering plant designed using k-medoids clustering.” Systematic biology 68, no. 4 (2019): 594–606.

Jónsson, Hákon, Aurélien Ginolhac, Mikkel Schubert, Philip LF Johnson, and Ludovic Orlando. “mapDamage2. 0: fast approximate Bayesian estimates of ancient DNA damage parameters.” Bioinformatics 29, no. 13 (2013): 1682–1684.

Jónsson, H. et al. (2014) ‘Speciation with gene flow in equids despite extensive chromosomal plasticity’, Proceedings of the National Academy of Sciences, 111(52), pp. 18655–18660. Available at: 10.1073/pnas.1412627111.

Kalyaanamoorthy, S. et al. (2017) ‘ModelFinder: fast model selection for accurate phylogenetic estimates’, Nature Methods, 14(6), pp. 587–589. Available at: 10.1038/nmeth.4285.

Katoh, K. and Standley, D.M. (2013) ‘MAFFT Multiple Sequence Alignment Software Version 7: Improvements in Performance and Usability’, Molecular Biology and Evolution, 30(4), pp. 772–780. Available at: 10.1093/molbev/mst010.

Keller, Andreas, Stephanie Kreis, Petra Leidinger, Frank Maixner, Nicole Ludwig, Christina Backes, Valentina Galata et al. “miRNAs in ancient tissue specimens of the Tyrolean Iceman.” Molecular Biology and Evolution 34, no. 4 (2017): 793–801.

Kim, D. et al. (2019) ‘Graph-based genome alignment and genotyping with HISAT2 and HISAT-genotype’, Nature Biotechnology, 37(8), pp. 907–915. Available at: 10.1038/s41587-019-0201-4.

Kourelis, J., et al. (2021) ‘RefPlantNLR is a comprehensive collection of experimentally validated plant disease resistance proteins from the NLR family’, PLOS Biology. Edited by X. Dong, 19(10), p. e3001124. Available at: 10.1371/journal.pbio.3001124.

Larson, D.A. et al. (2020) ‘A consensus phylogenomic approach highlights paleopolyploid and rapid radiation in the history of Ericales’, American Journal of Botany, 107(5), pp. 773–789. Available at: 10.1002/ajb2.1469.

Lagou, Loudmila Jelinscaia, Gudrun Kadereit, and Diego F. Morales-Briones. “Phylogenomic analysis of target enrichment and transcriptome data uncovers rapid radiation and extensive hybridization in the slipper orchid genus Cypripedium.” Annals of Botany 134, no. 7 (2024): 1229–1250.

Manni, M., et al. (2021) ‘BUSCO Update: Novel and Streamlined Workflows along with Broader and Deeper Phylogenetic Coverage for Scoring of Eukaryotic, Prokaryotic, and Viral Genomes’, Molecular Biology and Evolution. Edited by J. Kelley, 38(10), pp. 4647–4654. Available at: 10.1093/molbev/msab199.

Mármol-Sánchez, E. et al. (2023) ‘Historical RNA expression profiles from the extinct Tasmanian tiger’, Genome Research, 33(8), pp. 1299–1316. Available at: 10.1101/gr.277663.123.

Mármol-Sánchez, Emilio, Bastian Fromm, Nikolay Oskolkov, Zoé Pochon, Marianne Dehasque, Morteza Aslanzadeh, Elif Bozlak, et al. “Ancient RNA expression profiles from the extinct woolly mammoth.” Cell (2025).

Monteiro, F, and M. T. Nishimura. ‘Structural, functional, and genomic diversity of plant NLR proteins: an evolved resource for rational engineering of plant immunity.’ Annual review of phytopathology 56, no. 1 (2018): 243–267. Available at: 10.1146/annurev-phyto-080417-045817

Morales-Briones, D.F., et al. (2021) ‘Disentangling Sources of Gene Tree Discordance in Phylogenomic Data Sets: Testing Ancient Hybridizations in Amaranthaceae s.l’, Systematic Biology. Edited by R. Ree, 70(2), pp. 219–235. Available at: 10.1093/sysbio/syaa066.

Nguyen, L.-T. et al. (2015) ‘IQ-TREE: A Fast and Effective Stochastic Algorithm for Estimating Maximum-Likelihood Phylogenies’, Molecular Biology and Evolution, 32(1), pp. 268–274. Available at: 10.1093/molbev/msu300.

One Thousand Plant Transcriptomes Initiative. One thousand plant transcriptomes and the phylogenomics of green plants. Nature 574, 679–685 (2019).

Patro, R. et al. (2017) ‘Salmon provides fast and bias-aware quantification of transcript expression’, Nature Methods, 14(4), pp. 417–419. Available at: 10.1038/nmeth.4197.

R Core Team (2021). R: A language and environment for statistical computing. R Foundation for Statistical Computing, Vienna, Austria. URL https://www.R-project.org/.

Ramanauskas, K. and Igić, B. (2023) ‘kakapo : easy extraction and annotation of genes from raw RNA-seq reads’, PeerJ, 11, p. e16456. Available at: 10.7717/peerj.16456.

Rieux, A. et al. (2021) ‘Contribution of historical herbarium small RNAs to the reconstruction of a cassava mosaic geminivirus evolutionary history’. Sci. Rep. 11, 21280

Rollo, F. (1985) ‘Characterisation by molecular hybridization of RNA fragments isolated from ancient (1400 B.C.) seeds’, Theoretical and Applied Genetics, 71(2), pp. 330–333. Available at: 10.1007/BF00252076.

Rose, J.P. et al. (2018) ‘Phylogeny, historical biogeography, and diversification of angiosperm order Ericales suggest ancient Neotropical and East Asian connections’, Molecular Phylogenetics and Evolution, 122, pp. 59–79. Available at: 10.1016/j.ympev.2018.01.014.

Ruiz-Vargas, N. et al. (2024) ‘Transcriptome data from silica-preserved leaf tissue reveal gene flow patterns in a Caribbean bromeliad’, Annals of Botany, 133(3), pp. 459–472. Available at: 10.1093/aob/mcae002.

Sallon, S. et al. (2008) ‘Germination, Genetics, and Growth of an Ancient Date Seed’, Science, 320(5882), pp. 1464–1464. Available at: 10.1126/science.1153600.

Shaw, Barry, Carla L. Burrell, Darrell Green, Ana Navarro-Martinez, Daniel Scott, Anna Daroszewska, Rob van’t Hof, et al. “Molecular insights into an ancient form of Paget’s disease of bone.” Proceedings of the National Academy of Sciences 116, no. 21 (2019): 10463–10472.

Simão, Felipe A., Robert M. Waterhouse, Panagiotis Ioannidis, Evgenia V. Kriventseva, and Evgeny M. Zdobnov. “BUSCO: assessing genome assembly and annotation completeness with single-copy orthologs.” Bioinformatics 31, no. 19 (2015): 3210–3212.

Smith, O. et al. (2014) ‘Genomic methylation patterns in archaeological barley show de-methylation as a time-dependent diagenetic process’, Scientific Reports, 4(1), p. 5559. Available at: 10.1038/srep05559.

Smith, O. et al. Small RNA Activity in Archeological Barley Shows Novel Germination Inhibition in Response to Environment. Mol. Biol. Evol. 34, 2555–2562 (2017).

Smith, O., et al. (2019) ‘Ancient RNA from Late Pleistocene permafrost and historical canids shows tissue-specific transcriptome survival’, PLOS Biology. Edited by C. Tyler-Smith, 17(7), p. e3000166. Available at: 10.1371/journal.pbio.3000166.

Smith-Unna, R. et al. (2016) ‘TransRate: reference-free quality assessment of de novo transcriptome assemblies’, Genome Research, 26(8), pp. 1134–1144. Available at: 10.1101/gr.196469.115.

Song, L. and Florea, L. (2015) ‘Rcorrector: efficient and accurate error correction for Illumina RNA-seq reads’, Gigascience, 4(1), pp. s13742–015-0089-y. Available at: 10.1186/s13742-015-0089-y.

Speer, Kelly A., Melissa TR Hawkins, Mary Faith C. Flores, Michael R. McGowen, Robert C. Fleischer, Jesús E. Maldonado, Michael G. Campana, and Carly R. Muletz-Wolz. “A comparative study of RNA yields from museum specimens, including an optimized protocol for extracting RNA from formalin-fixed specimens.” Frontiers in Ecology and Evolution 10 (2022): 953131.

Tyszka, A.S., Larson, D.A. and Walker, J.F. (2025) ‘Sequencing historical RNA: unrealized potential to increase understanding of the plant tree of life’, Trends in Plant Science, p. S1360138524003054. Available at: 10.1016/j.tplants.2024.11.004.

Van Dongen, S. (2008) ‘Graph Clustering Via a Discrete Uncoupling Process’, SIAM Journal on Matrix Analysis and Applications, 30(1), pp. 121–141. Available at: 10.1137/040608635.

Venables, William N., and Brian D. Ripley. Modern applied statistics with S. Springer Science & Business Media, 2013.

Venanzi, F. M. & Rollo, F. ‘Mummy RNA lasts longer’. Nature 343, 25–26 (1990).

Walker, J.F., et al. (2022) ‘Concordance-Based Approaches for the Inference of Relationships and Molecular Rates with Phylogenomic Data Sets’, Systematic Biology. Edited by R. Thomson, 71(4), pp. 943–958. Available at: 10.1093/sysbio/syab052.

Wong, Darren CJ, and Rod Peakall. “Orchid phylotranscriptomics: the prospects of repurposing multi-tissue transcriptomes for phylogenetic analysis and beyond.” Frontiers in Plant Science 13 (2022): 910362.

Wood, D.E., Lu, J. and Langmead, B. (2019) ‘Improved metagenomic analysis with Kraken 2’, Genome Biology, 20(1), p. 257. Available at: 10.1186/s13059-019-1891-0.

Woodcock-Girard, Miles D., Eric C. Bretz, Holly M. Robertson, Karolis Ramanauskas, Jarrad T. Hampton-Marcell, and Joseph F. Walker. “Semblans: automated assembly and processing of RNA-seq data.” Bioinformatics 41, no. 1 (2025): btaf003.

Xie, J. et al. (2023) ‘Tree Visualization By One Table (tvBOT): a web application for visualizing, modifying and annotating phylogenetic trees’, Nucleic Acids Research, 51(W1), pp. W587–W592. Available at: 10.1093/nar/gkad359.

Yang, Y. and Smith, S.A. (2013) ‘Optimizing de novo assembly of short-read RNA-seq data for phylogenomics’, BMC Genomics, 14(1), p. 328. Available at: 10.1186/1471-2164-14-328.

Yang, Ya, and Stephen A. Smith. “Orthology inference in nonmodel organisms using transcriptomes and low-coverage genomes: improving accuracy and matrix occupancy for phylogenomics.” Molecular biology and evolution 31, no. 11 (2014): 3081–3092.

Zhang, C. et al. (2018) ‘ASTRAL-III: polynomial time species tree reconstruction from partially resolved gene trees’, BMC Bioinformatics, 19(S6), p. 153. Available at: 10.1186/s12859-018-2129-y.

Zuntini, A.R. et al. (2024) ‘Phylogenomics and the rise of the angiosperms’, Nature, 629(8013), pp. 843–850. Available at: 10.1038/s41586-024-07324-0.

