## Supplemental Fig. 1 for "Herbaria provide a valuable resource for obtaining informative mRNA"

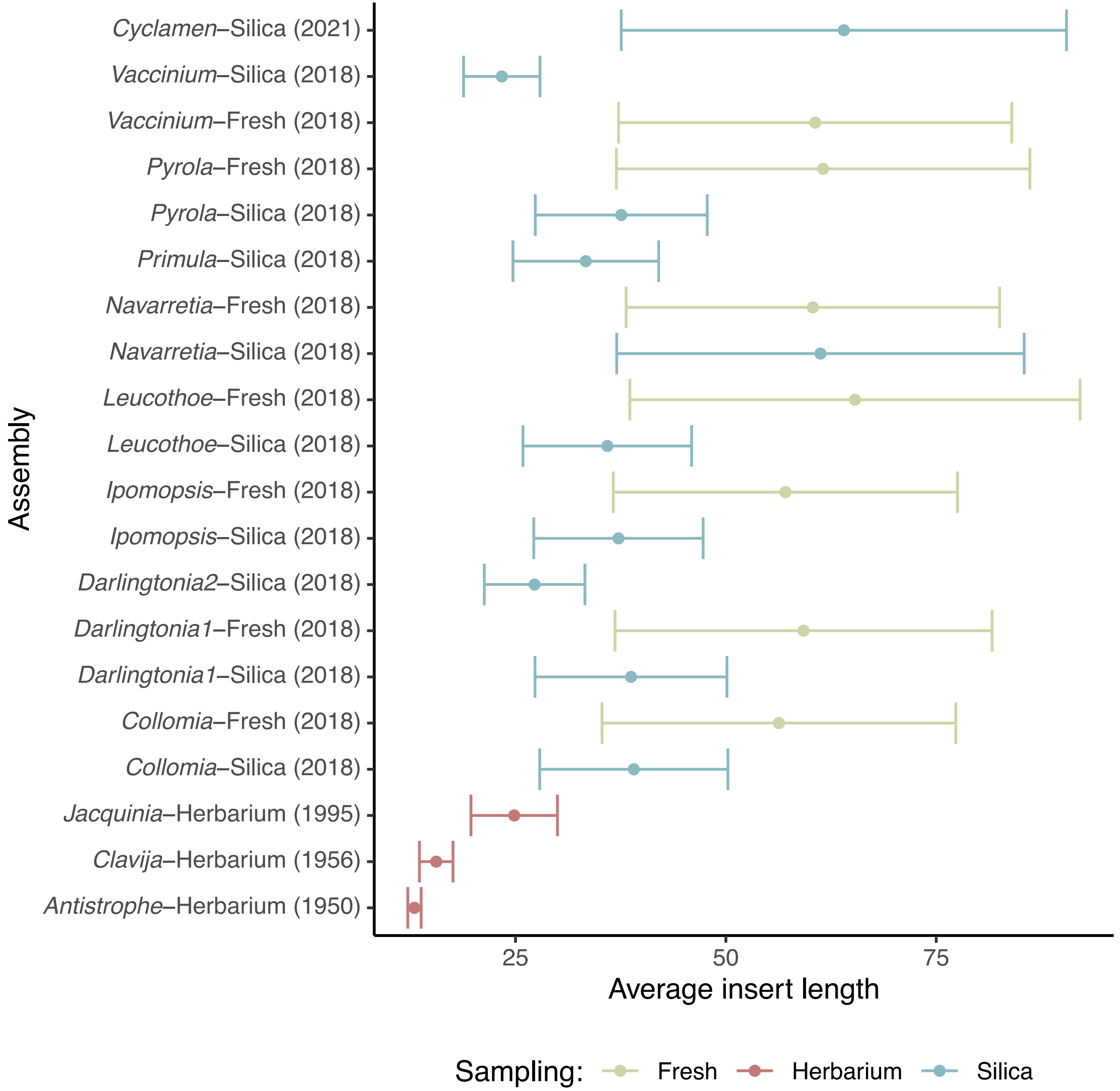


**Supplemental Figure 1:** Average and standard deviation insert length for each assembly, not including zero length inserts.
