## Supplemental Fig. 2 for "Herbaria provide a valuable resource for obtaining informative mRNA"

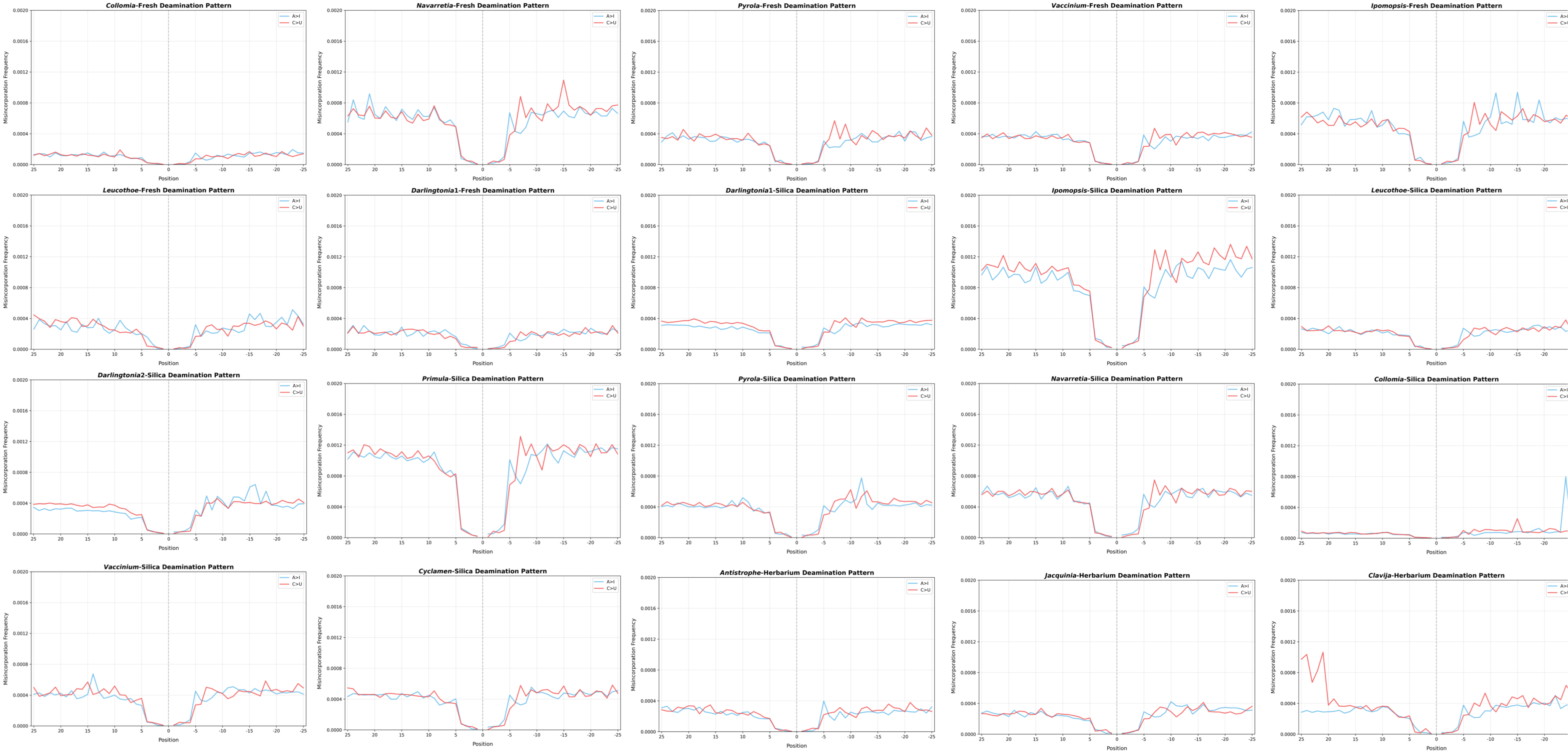


**Supplemental Figure 2:** Deamination adenosine to inosine (A>I) and cytidine to uridine (C>U) tested for all samples.
