## Supplemental Fig. 3 for "Herbaria provide a valuable resource for obtaining informative mRNA"

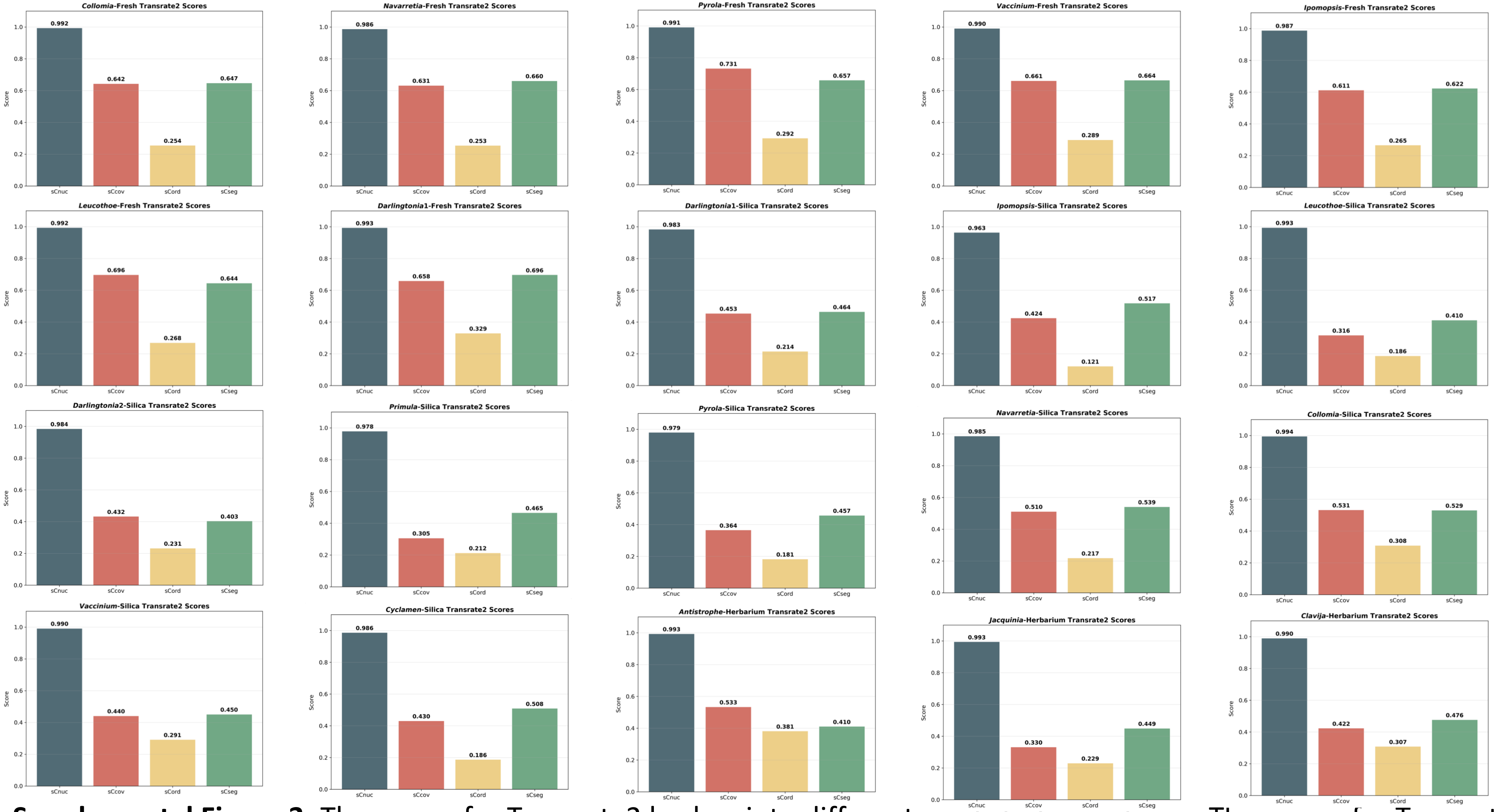


**Supplemental Figure 3:** The scores for Transrate2 broken into different predicted error types: The scores for Transrate2 broken into different predicted error types: reliability of the base pair calls (*sCnuc*), the coverage level of the transcript (*sCov*), the pairing of the mapped reads (*sCord*); and evidence of chimerism within the transcripts (*sCseg*). Can you look over the wording on this, not sure the best way to do about it.
