## Supplemental Fig. 4 for "Herbaria provide a valuable resource for obtaining informative mRNA"

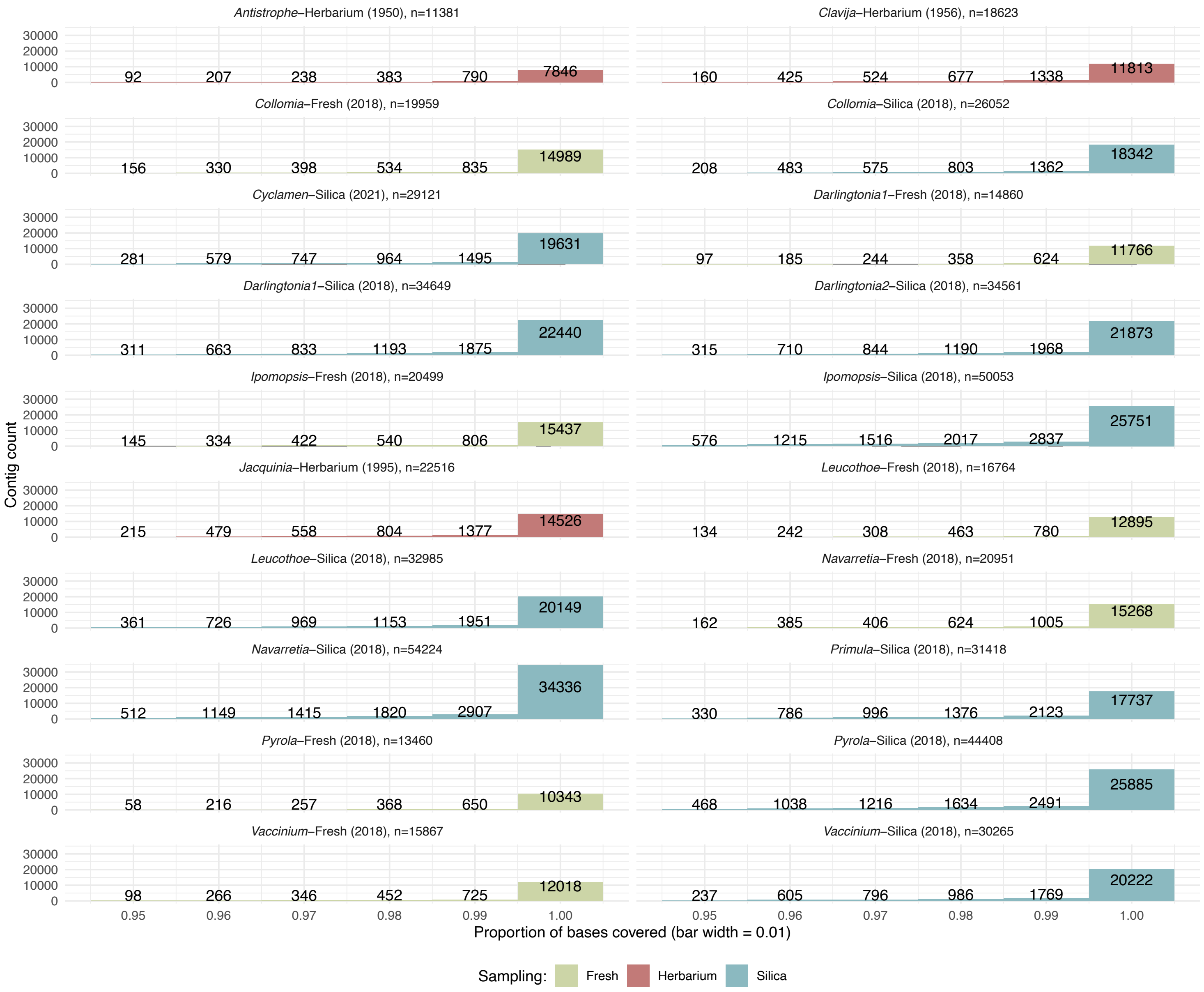


**Supplemental Figure 4:** The assessed coverage for each transcriptome on a contig-by-contig basis, displaying contigs with at least 95% coverage.
