## Supplemental Fig. 5 for "Herbaria provide a valuable resource for obtaining informative mRNA"

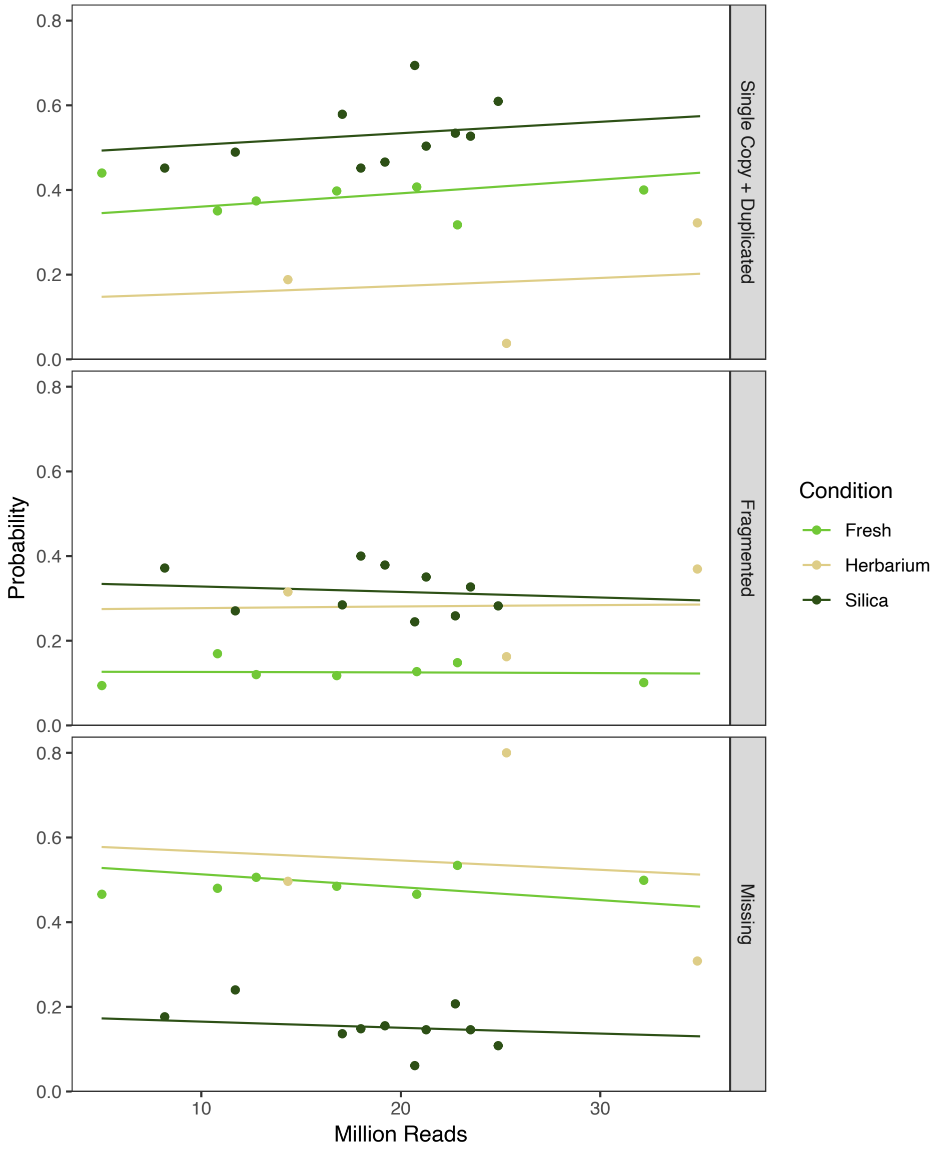


**Supplemental Figure 5:** Recovery of BUSCO loci in transcriptome assemblies from different storage conditions. Multinomial logistic regression of BUSCO recovery (complete = single-copy + duplicated, fragmented, and missing) against storage condition and sequencing depth (in millions of reads). Points give the observed proportions per sample, while lines are the predicted probability of each category over the range of read depth, conditioning on the storage condition.
