## Supplemental Document 1 for "Herbaria provide a valuable resource for obtaining informative mRNA"

### Ipomopsis-Silica

**B1: 3**

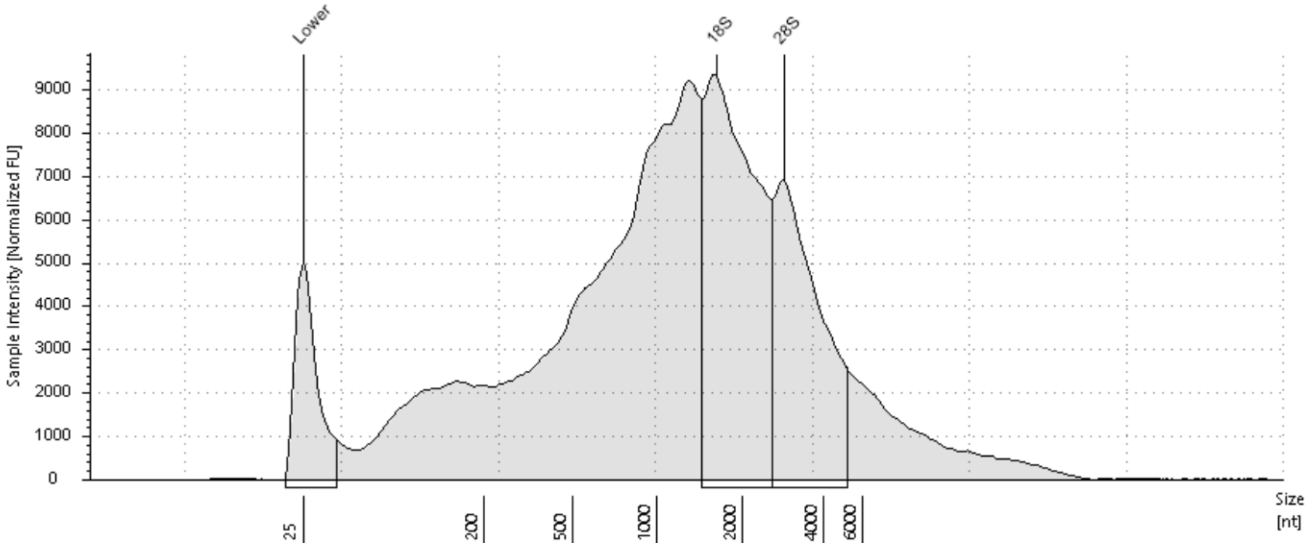


**Sample Table**

| Well | RINe | 28S/18S (Area) | Conc. [ng/µl] | Sample Description | Alert | Observations |
|---|---|---|---|---|---|---|
| B1 | 3.2 | 0.7 | 366 | 3 | | |

**Peak Table**

| Size [nt] | Calibrated Conc. [ng/µl] | Assigned Conc. [ng/µl] | Peak Molarity [nmol/l] | % Integrated Area | Peak Comment | Observations |
|---|---|---|---|---|---|---|
| 25 | 40.0 | 40.0 | 4710 | - | | Lower Marker |
| 1622 | 83.5 | - | 151 | 60.45 | | 18S |
| 2852 | 54.6 | - | 56.3 | 39.55 | | 28S |

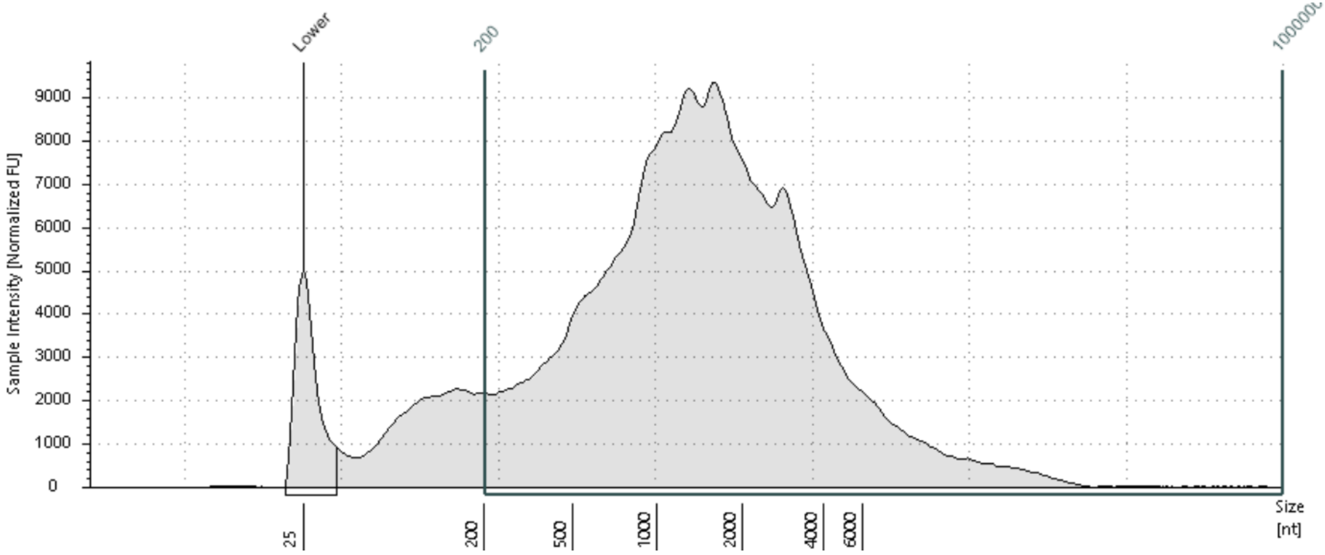


**Region Table**

| From [nt] | To [nt] | Average Size [nt] | Conc. [ng/µl] | Region Molarity [nmol/l] | % of Total | Region Comment | Color |
|---|---|---|---|---|---|---|---|
| 200 | 1000000 | 12962 | 331 | 75.0 | 90.46 | DV200 | |

### Leucothoe-Silica

**D1: 7**

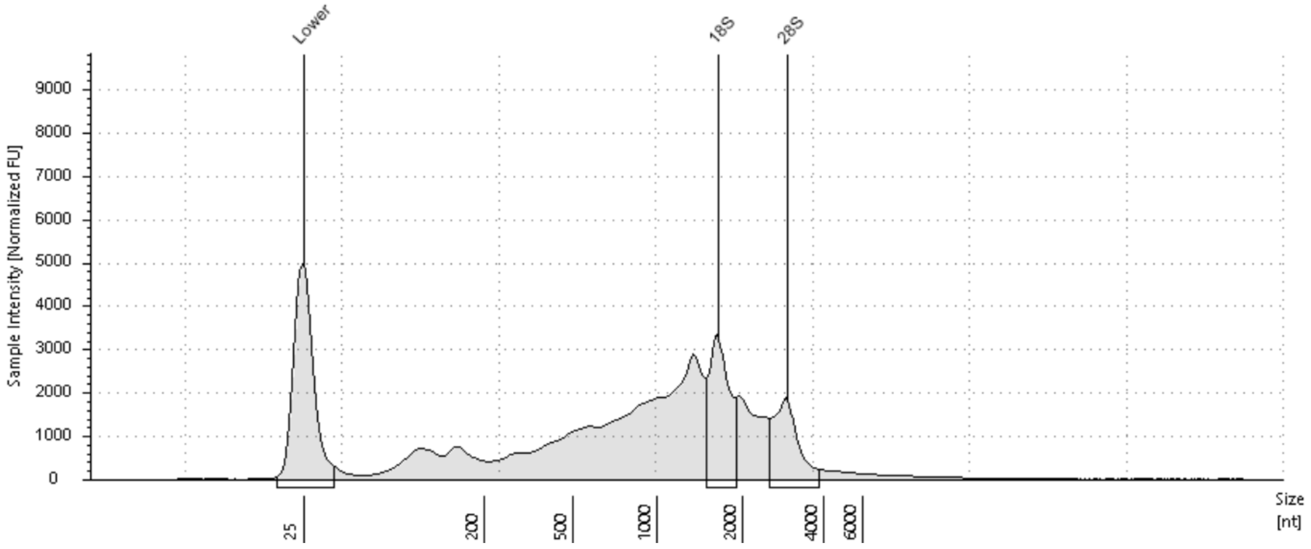


##### Sample Table

| Well | RINe | 28S/18S (Area) | Conc. [ng/µl] | Sample Description | Alert | Observations |
|---|---|---|---|---|---|---|
| D1 | 4.2 | 0.7 | 91.8 | 7 | | |

##### Peak Table

| Size [nt] | Calibrated Conc. [ng/µl] | Assigned Conc. [ng/µl] | Peak Molarity [nmol/l] | % Integrated Area | Peak Comment | Observations |
|---|---|---|---|---|---|---|
| 25 | 40.0 | 40.0 | 4710 | - | | Lower Marker |
| 1647 | 12.8 | - | 22.9 | 59.46 | | 18S |
| 2944 | 8.76 | - | 8.75 | 40.54 | | 28S |

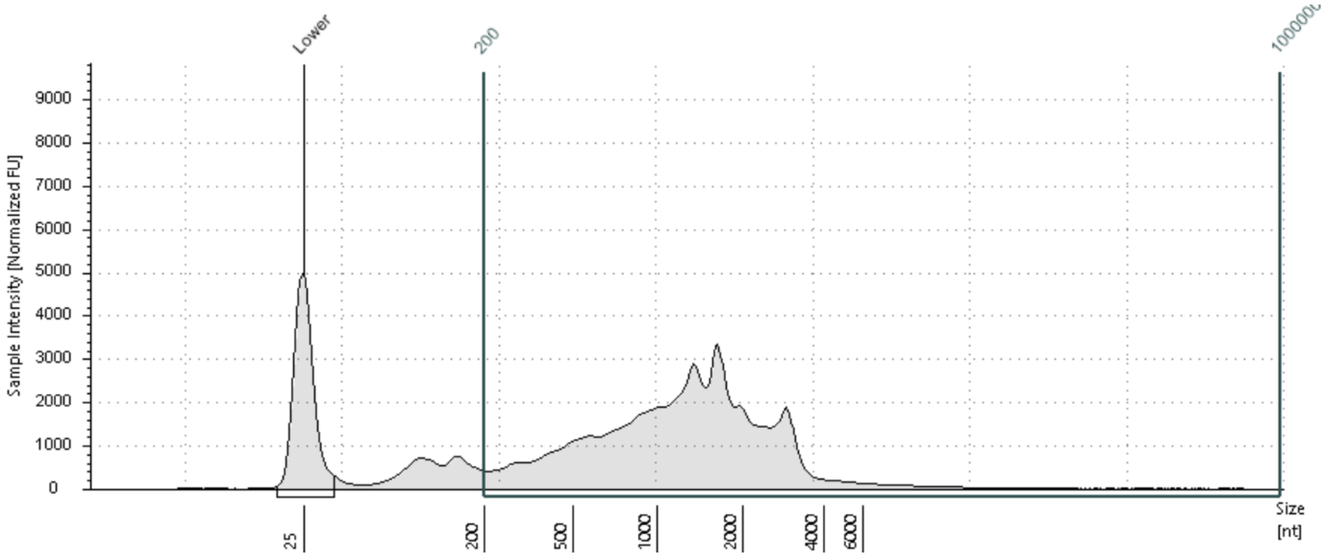


##### Region Table

| From [nt] | To [nt] | Average Size [nt] | Conc. [ng/µl] | Region Molarity [nmol/l] | % of Total | Region Comment | Color |
|---|---|---|---|---|---|---|---|
| 200 | 1000000 | 4929 | 81.6 | 48.7 | 88.89 | DV200 | |


© Agilent Technologies, Inc. 2021

### Darlingtonia2-Silica

**E1: 8**

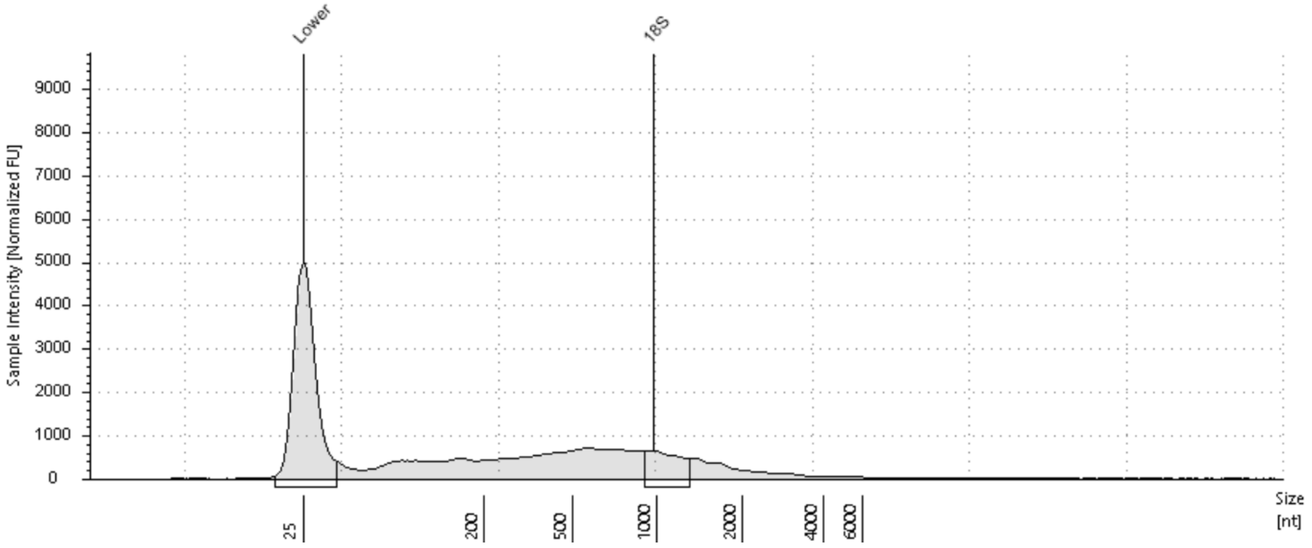


#### Sample Table

| Well | RINe | 28S/18S (Area) | Conc. [ng/µl] | Sample Description | Alert | Observations |
|---|---|---|---|---|---|---|
| E1 | 2.7 | - | 30.4 | 8 | | The upper ribosomal fragment has degraded |

#### Peak Table

| Size [nt] | Calibrated Conc. [ng/µl] | Assigned Conc. [ng/µl] | Peak Molarity [nmol/l] | % Integrated Area | Peak Comment | Observations |
|---|---|---|---|---|---|---|
| 25 | 40.0 | 40.0 | 4710 | - | | Lower Marker |
| 972 | 3.84 | - | 11.6 | 100.00 | | 18S |

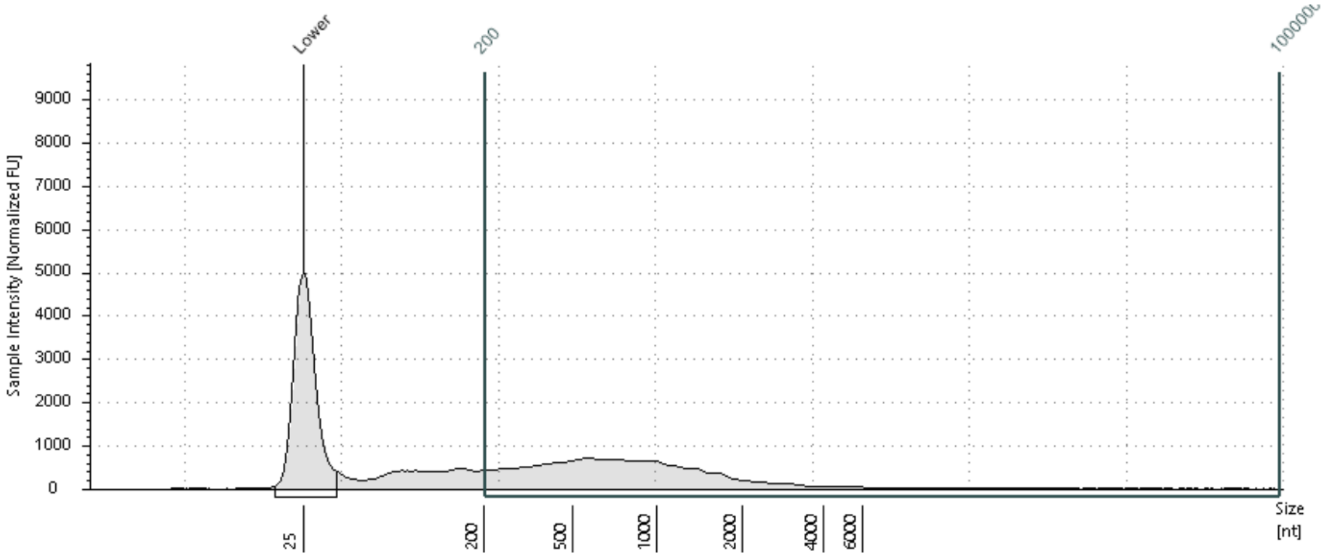


#### Region Table

| From [nt] | To [nt] | Average Size [nt] | Conc. [ng/µl] | Region Molarity [nmol/l] | % of Total | Region Comment | Color |
|---|---|---|---|---|---|---|---|
| 200 | 1000000 | 9123 | 22.5 | 7.24 | 73.83 | DV200 | |

 © Agilent Technologies, Inc. 2021 

### Primula-Silica

**F1: 9**

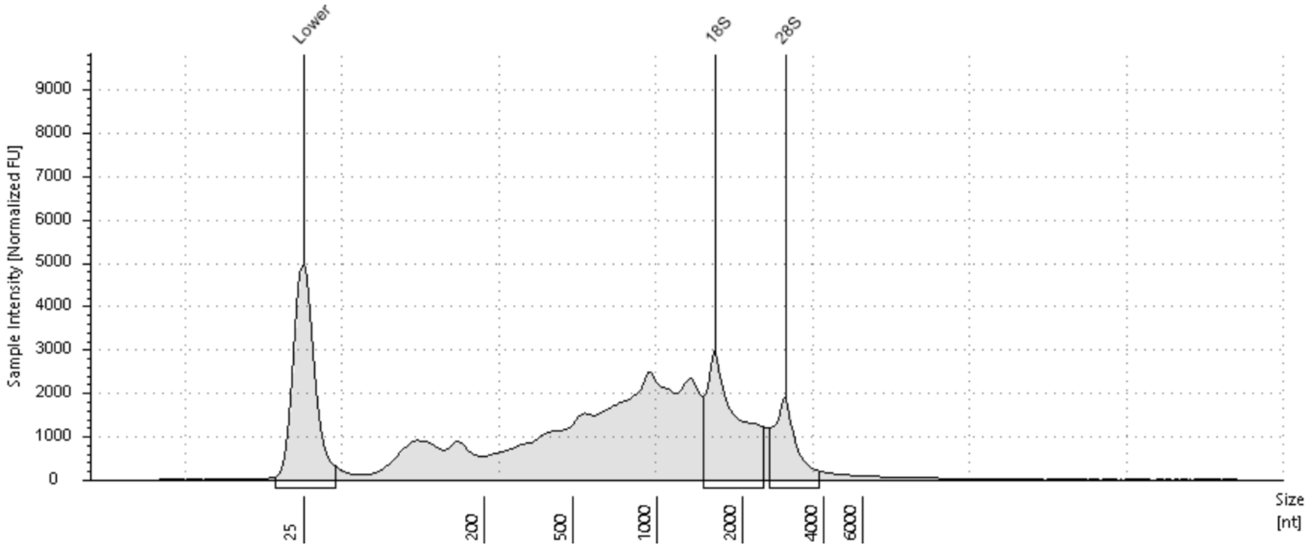


**Sample Table**

| Well | RINe | 28S/18S (Area) | Conc. [ng/µl] | Sample Description | Alert | Observations |
|---|---|---|---|---|---|---|
| F1 | 3.6 | 0.4 | 89.4 | 9 | | |

**Peak Table**

| Size [nt] | Calibrated Conc. [ng/µl] | Assigned Conc. [ng/µl] | Peak Molarity [nmol/l] | % Integrated Area | Peak Comment | Observations |
|---|---|---|---|---|---|---|
| 25 | 40.0 | 40.0 | 4710 | - | | Lower Marker |
| 1604 | 16.7 | - | 30.6 | 69.34 | | 18S |
| 2887 | 7.38 | - | 7.52 | 30.66 | | 28S |

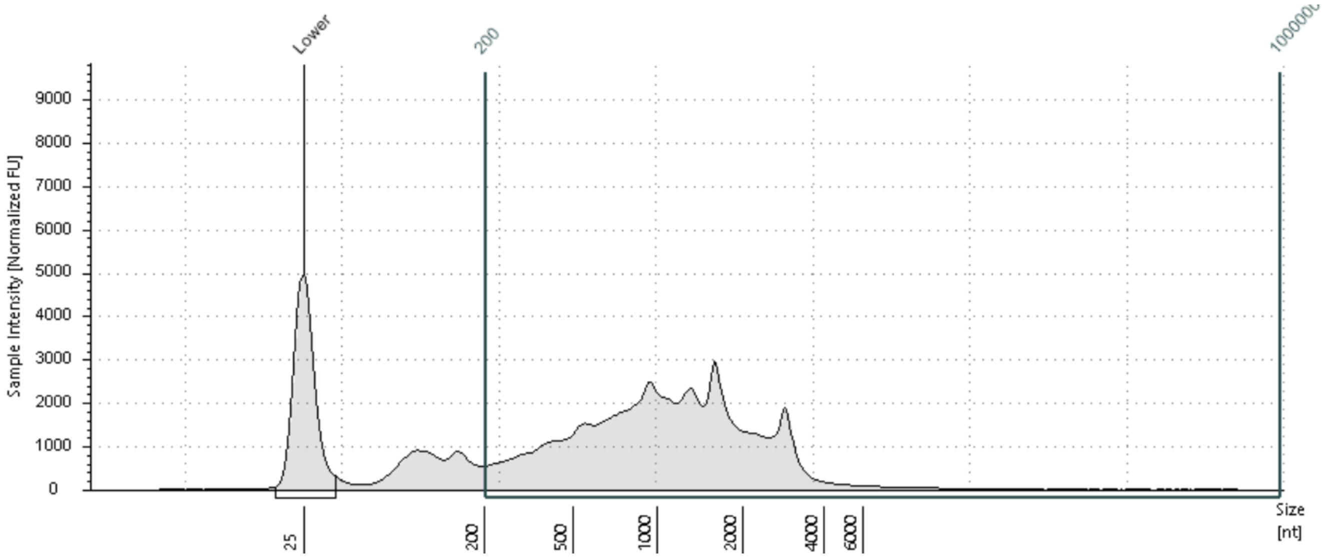


**Region Table**

| From [nt] | To [nt] | Average Size [nt] | Conc. [ng/µl] | Region Molarity [nmol/l] | % of Total | Region Comment | Color |
|---|---|---|---|---|---|---|---|
| 200 | 1000000 | 3816 | 77.1 | 59.5 | 86.26 | DV200 | ■ |


© Agilent Technologies, Inc. 2021

### Pyrola-Silica

**G1: 11**

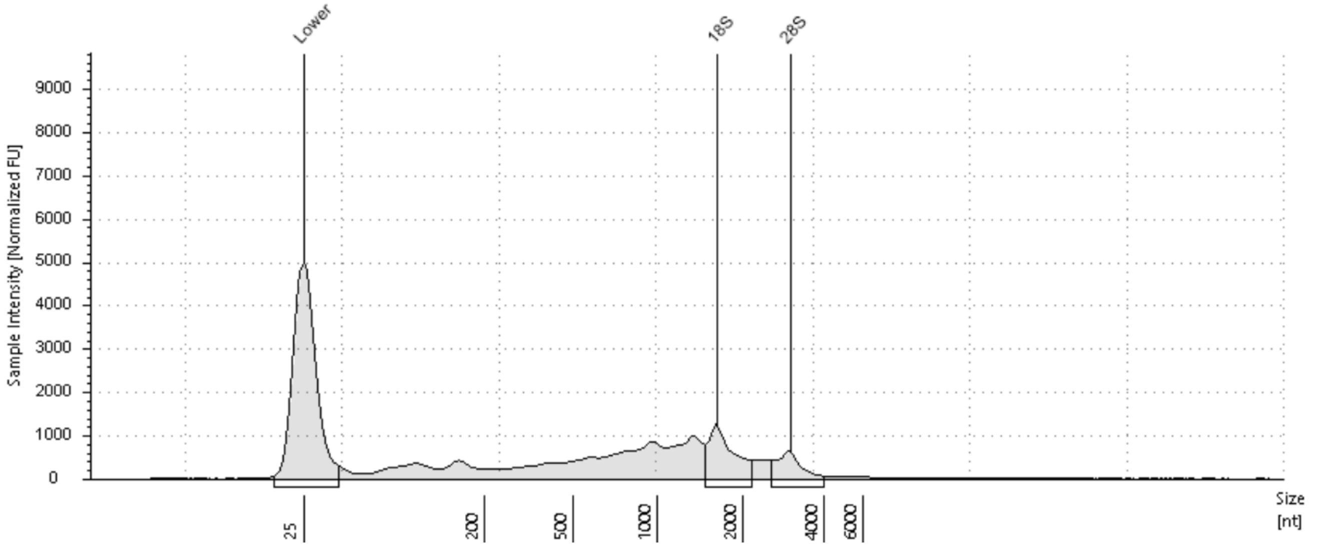


**Sample Table**

| Well | RINe | 28S/18S (Area) | Conc. [ng/µl] | Sample Description | Alert | Observations |
|---|---|---|---|---|---|---|
| G1 | 4.1 | 0.5 | 31.0 | 11 | | |

**Peak Table**

| Size [nt] | Calibrated Conc. [ng/µl] | Assigned Conc. [ng/µl] | Peak Molarity [nmol/l] | % Integrated Area | Peak Comment | Observations |
|---|---|---|---|---|---|---|
| 25 | 40.0 | 40.0 | 4710 | - | | Lower Marker |
| 1624 | 5.01 | - | 9.06 | 66.19 | | 18S |
| 2995 | 2.56 | - | 2.51 | 33.81 | | 28S |

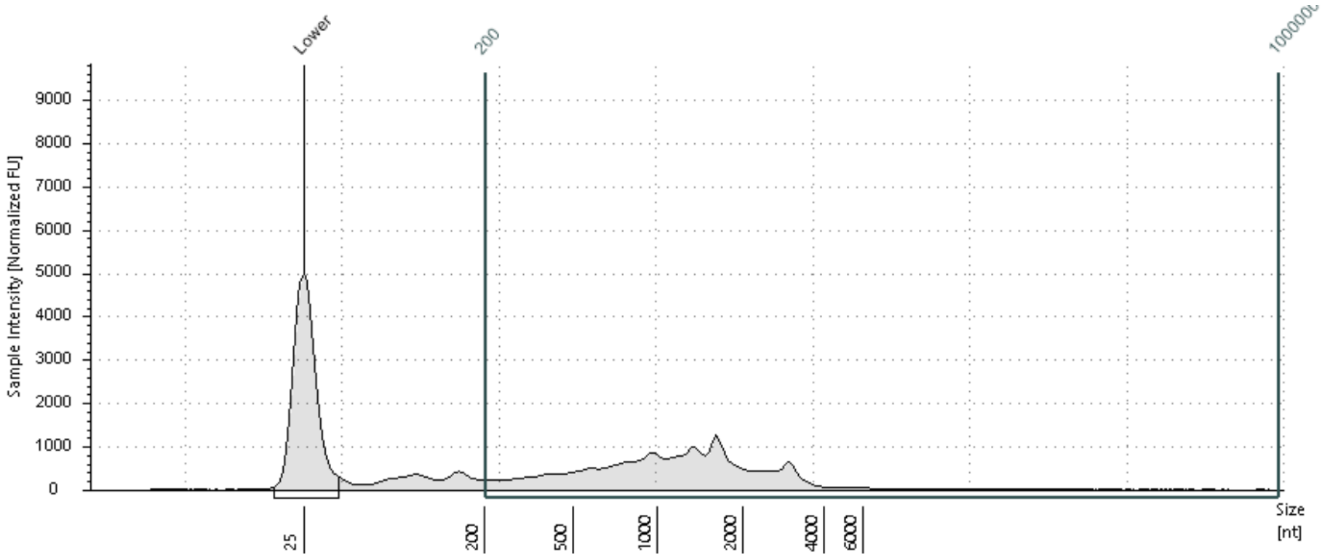


**Region Table**

| From [nt] | To [nt] | Average Size [nt] | Conc. [ng/µl] | Region Molarity [nmol/l] | % of Total | Region Comment | Color |
|---|---|---|---|---|---|---|---|
| 200 | 1000000 | 5990 | 25.7 | 12.6 | 82.80 | DV200 | |

 © Agilent Technologies, Inc. 2021 

### Navarretia-Silica

**H1: 12**

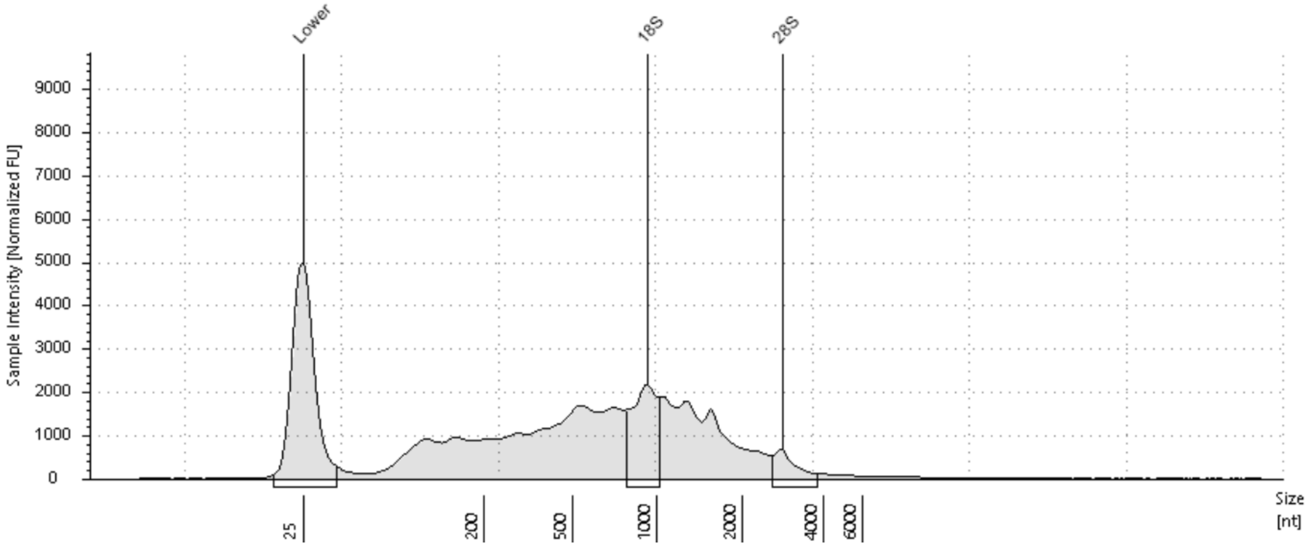


**Sample Table**

| Well | RINe | 28S/18S (Area) | Conc. [ng/µl] | Sample Description | Alert | Observations |
|---|---|---|---|---|---|---|
| H1 | 3.7 | 0.3 | 71.5 | 12 | | The upper ribosomal fragment has degraded; RINe edited |

**Peak Table**

| Size [nt] | Calibrated Conc. [ng/µl] | Assigned Conc. [ng/µl] | Peak Molarity [nmol/l] | % Integrated Area | Peak Comment | Observations |
|---|---|---|---|---|---|---|
| 25 | 40.0 | 40.0 | 4710 | - | | Lower Marker |
| 924 | 9.33 | - | 29.7 | 78.74 | | 18S edited |
| 2812 | 2.52 | - | 2.63 | 21.26 | | 28S |

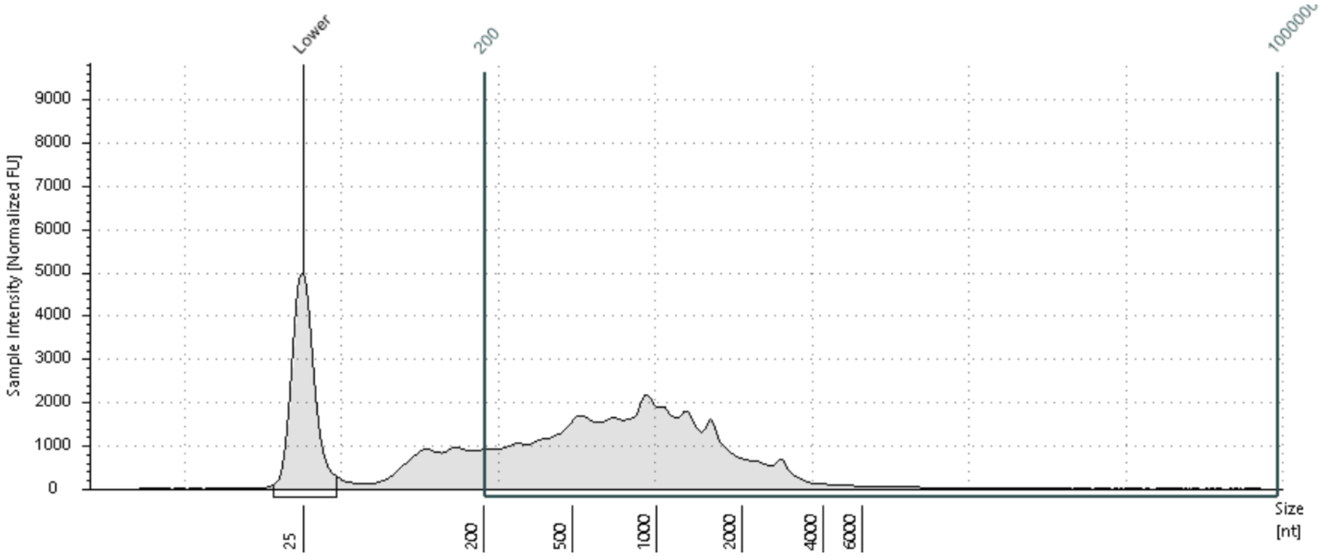


**Region Table**

| From [nt] | To [nt] | Average Size [nt] | Conc. [ng/µl] | Region Molarity [nmol/l] | % of Total | Region Comment | Color |
|---|---|---|---|---|---|---|---|
| 200 | 1000000 | 4929 | 59.3 | 35.4 | 82.90 | DV200 | |

 © Agilent Technologies, Inc. 2021 

### Collomia-Silica

**A2: 13**

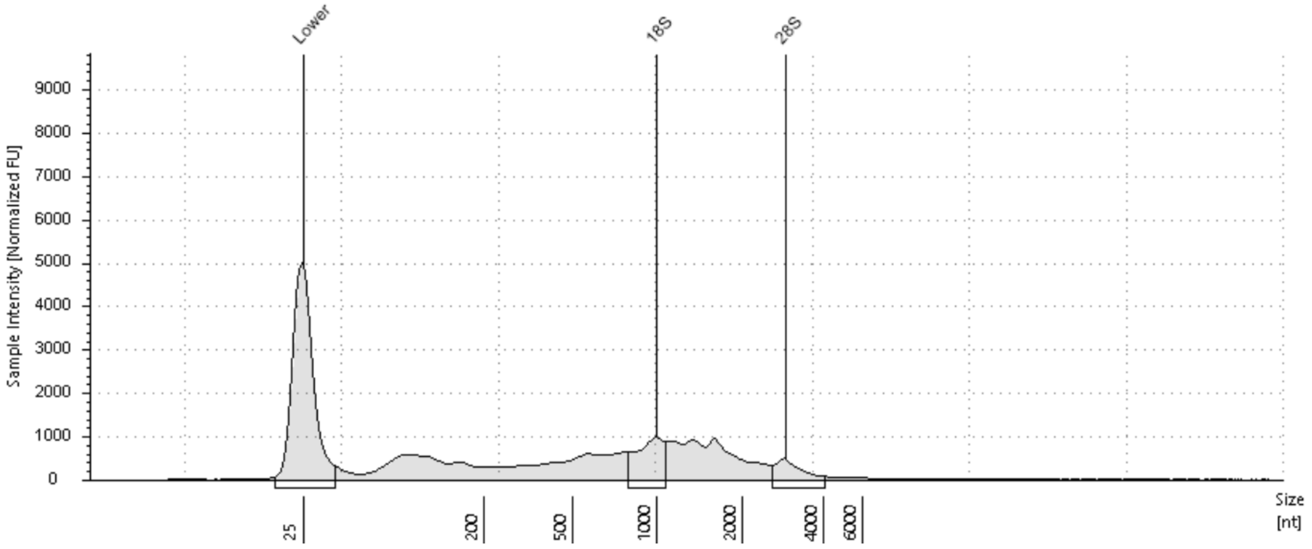


**Sample Table**

| Well | RINe | 28S/18S (Area) | Conc. [ng/µl] | Sample Description | Alert | Observations |
|---|---|---|---|---|---|---|
| A2 | 4.3 | 0.5 | 37.7 | 13 | | The upper ribosomal fragment has degraded |

**Peak Table**

| Size [nt] | Calibrated Conc. [ng/µl] | Assigned Conc. [ng/µl] | Peak Molarity [nmol/l] | % Integrated Area | Peak Comment | Observations |
|---|---|---|---|---|---|---|
| 25 | 40.0 | 40.0 | 4710 | - | | Lower Marker |
| 998 | 4.74 | - | 14.0 | 67.91 | | 18S |
| 2892 | 2.24 | - | 2.28 | 32.09 | | 28S |

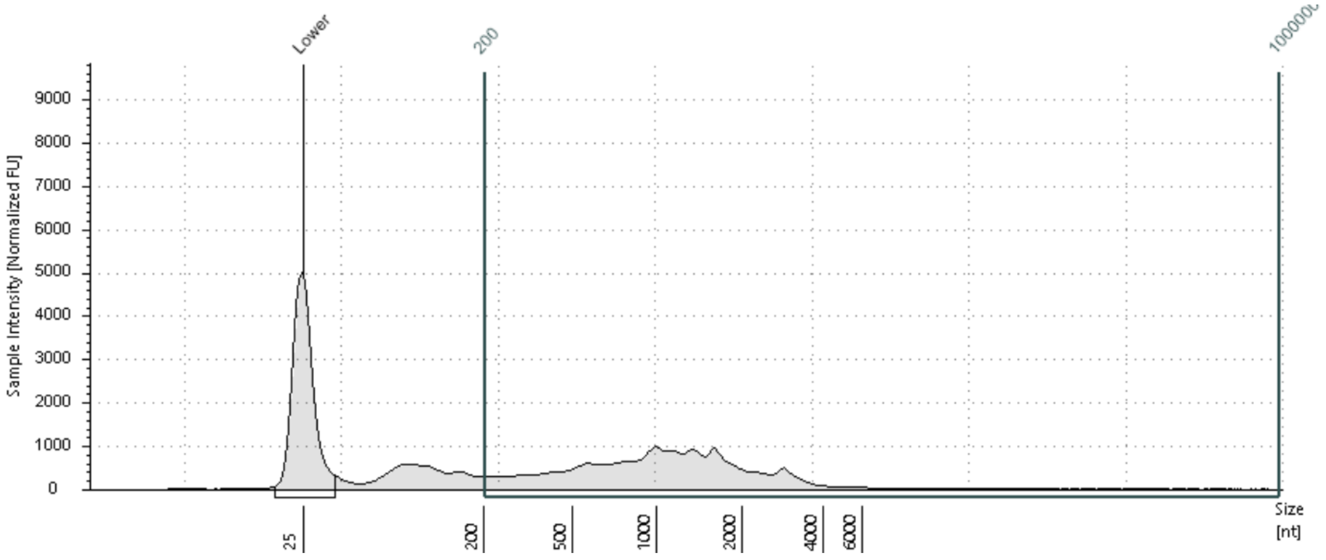


**Region Table**

| From [nt] | To [nt] | Average Size [nt] | Conc. [ng/µl] | Region Molarity [nmol/l] | % of Total | Region Comment | Color |
|---|---|---|---|---|---|---|---|
| 200 | 1000000 | 6038 | 29.1 | 14.2 | 77.16 | DV200 | |

 © Agilent Technologies, Inc. 2021 

### Vaccinium-Silica

**B2: 14**

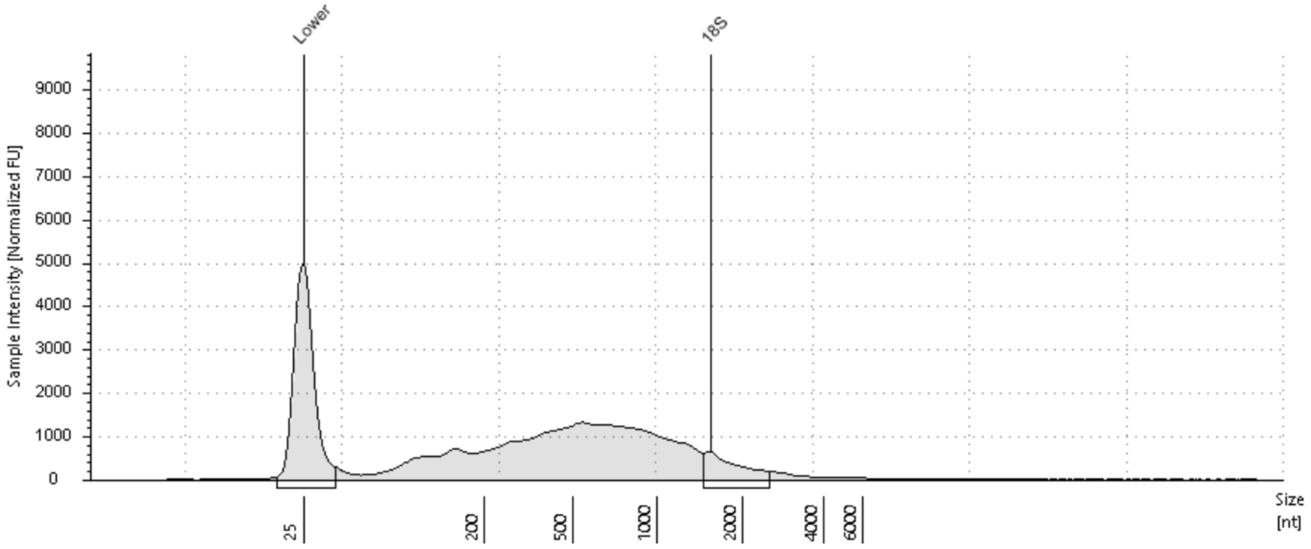


**Sample Table**

| Well | RINe | 28S/18S (Area) | Conc. [ng/µl] | Sample Description | Alert | Observations |
|---|---|---|---|---|---|---|
| B2 | 1.8 | - | 53.0 | 14 | | The upper ribosomal fragment has degraded |

**Peak Table**

| Size [nt] | Calibrated Conc. [ng/µl] | Assigned Conc. [ng/µl] | Peak Molarity [nmol/l] | % Integrated Area | Peak Comment | Observations |
|---|---|---|---|---|---|---|
| 25 | 40.0 | 40.0 | 4710 | - | | Lower Marker |
| 1557 | 4.15 | - | 7.84 | 100.00 | | 18S |

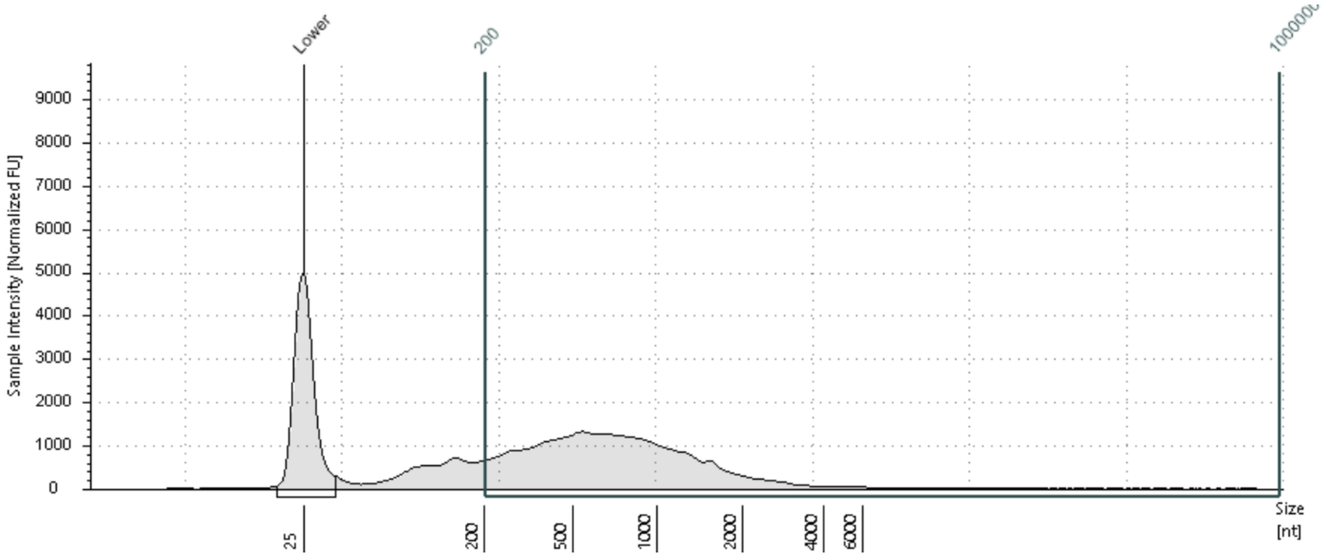


**Region Table**

| From [nt] | To [nt] | Average Size [nt] | Conc. [ng/µl] | Region Molarity [nmol/l] | % of Total | Region Comment | Color |
|---|---|---|---|---|---|---|---|
| 200 | 1000000 | 4485 | 43.2 | 28.3 | 81.43 | DV200 | |

### Cyclamen-Silica

## C2: 18

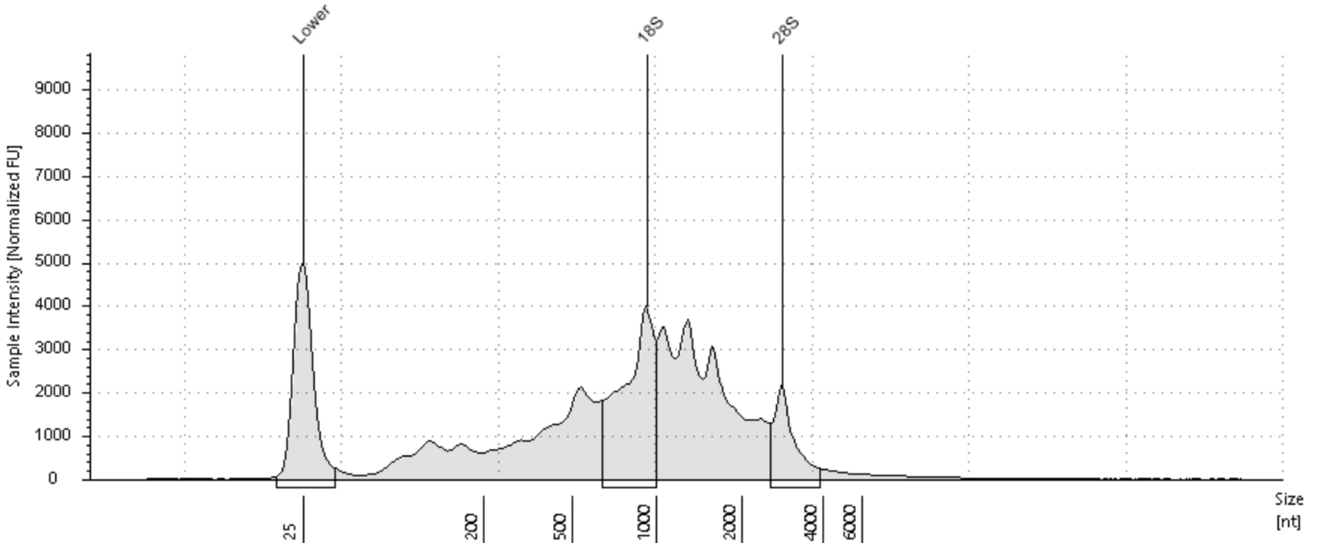


##### Sample Table

| Well | RINe | 28S/18S (Area) | Conc. [ng/µl] | Sample Description | Alert | Observations |
|---|---|---|---|---|---|---|
| C2 | 5.0 | 0.4 | 115 | 18 | | |

##### Peak Table

| Size [nt] | Calibrated Conc. [ng/µl] | Assigned Conc. [ng/µl] | Peak Molarity [nmol/l] | % Integrated Area | Peak Comment | Observations |
|---|---|---|---|---|---|---|
| 25 | 40.0 | 40.0 | 4710 | - | | Lower Marker |
| 930 | 24.1 | - | 76.2 | 74.07 | | 18S |
| 2834 | 8.44 | - | 8.76 | 25.93 | | 28S |

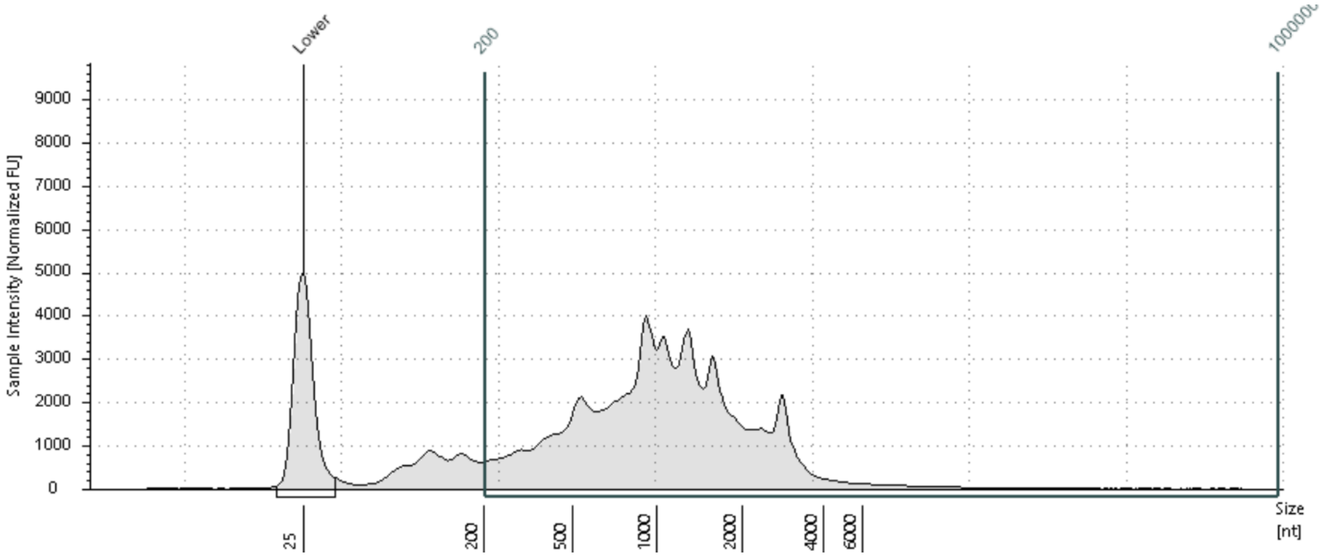


##### Region Table

| From [nt] | To [nt] | Average Size [nt] | Conc. [ng/µl] | Region Molarity [nmol/l] | % of Total | Region Comment | Color |
|---|---|---|---|---|---|---|---|
| 200 | 1000000 | 4601 | 103 | 65.8 | 89.66 | DV200 | |

 © Agilent Technologies, Inc. 2021 

### Darlingtonia1-Silica

**A1: 2**

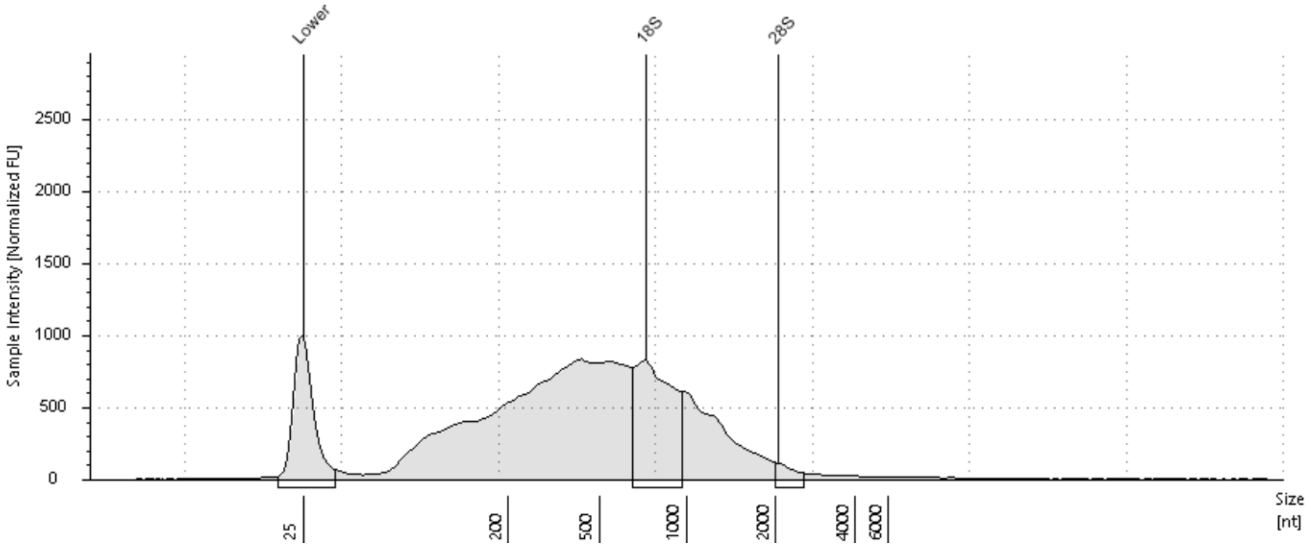


**Sample Table**

| Well | RINe | 28S/18S (Area) | Conc. [pg/µl] | Sample Description | Alert | Observations |
|---|---|---|---|---|---|---|
| A1 | 2.4 | 0.1 | 2550 | 2 | | The upper ribosomal fragment has degraded |

**Peak Table**

| Size [nt] | Calibrated Conc. [pg/µl] | Assigned Conc. [pg/µl] | Peak Molarity [pmol/l] | % Integrated Area | Peak Comment | Observations |
|---|---|---|---|---|---|---|
| 25 | 700 | 700 | 82400 | - | | Lower Marker |
| 726 | 451 | - | 1830 | 94.00 | | 18S |
| 2053 | 28.8 | - | 41.2 | 6.00 | | 28S |

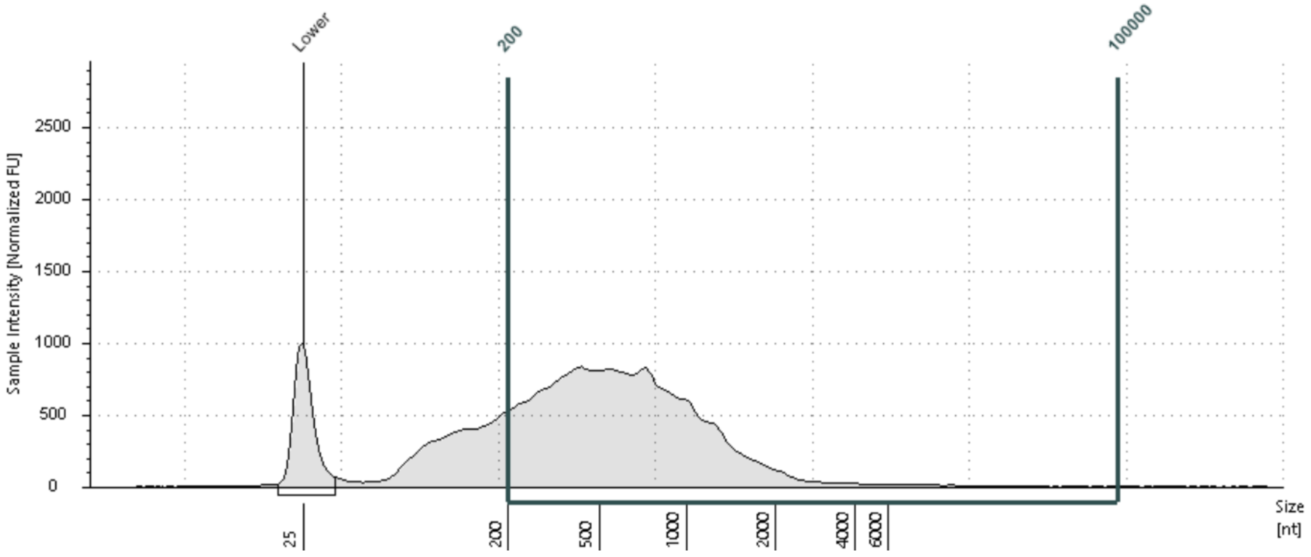


**Region Table**

| From [nt] | To [nt] | Average Size [nt] | Conc. [pg/µl] | Region Molarity [pmol/l] | % of Total | Region Comment | Color |
|---|---|---|---|---|---|---|---|
| 200 | 100000 | 4683 | 2040 | 1280 | 79.93 | DV200 | |

 © Agilent Technologies, Inc. 2021 

### Jacquinia-Herbarium

**D1: 25**

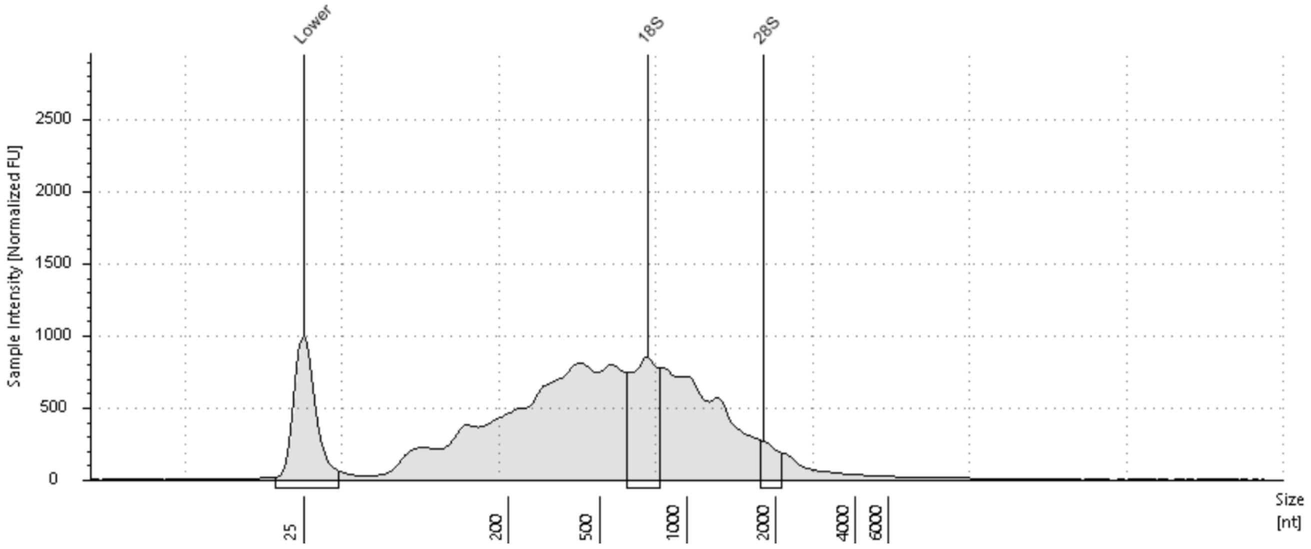


**Sample Table**

| Well | RINe | 28S/18S (Area) | Conc. [pg/µl] | Sample Description | Alert | Observations |
|---|---|---|---|---|---|---|
| D1 | 2.6 | 0.2 | 2380 | 25 | | The upper ribosomal fragment has degraded |

**Peak Table**

| Size [nt] | Calibrated Conc. [pg/µl] | Assigned Conc. [pg/µl] | Peak Molarity [pmol/l] | % Integrated Area | Peak Comment | Observations |
|---|---|---|---|---|---|---|
| 25 | 700 | 700 | 82400 | - | | Lower Marker |
| 730 | 304 | - | 1220 | 83.46 | | 18S |
| 1811 | 60.3 | - | 97.8 | 16.54 | | 28S |

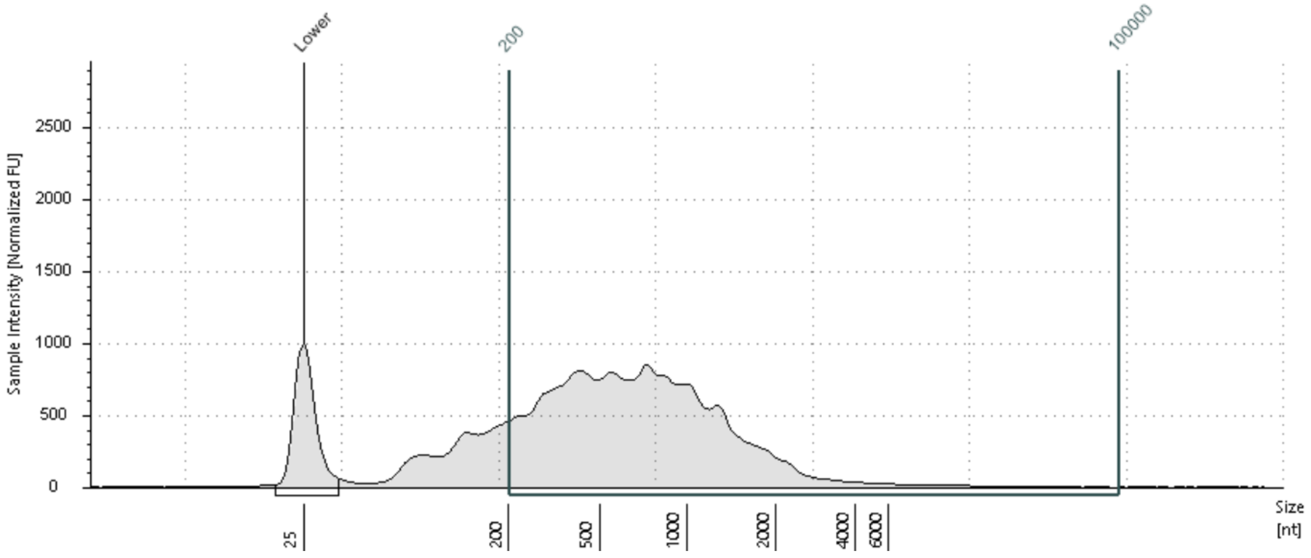


**Region Table**

| From [nt] | To [nt] | Average Size [nt] | Conc. [pg/µl] | Region Molarity [pmol/l] | % of Total | Region Comment | Color |
|---|---|---|---|---|---|---|---|
| 200 | 100000 | 6250 | 1980 | 932 | 83.29 | DV200 | |

 © Agilent Technologies, Inc. 2021 
